# Mice Mucosal Leishmaniasis model shown high parasite load, increased cytotoxicity and impaired IL-10^+^ cells response

**DOI:** 10.1101/2025.04.22.650112

**Authors:** Alisson Amaral da Rocha, Júlio Souza dos Santos, Igor Bittencourt dos Santos, Douglas B. de Almeida, Naiara Carla dos Santos Manhães, Hozany Praxedes dos Santos, João Victor Paiva Romano, Elias Barbosa da Silva Junior, Antônio José da Silva Gonçalves, Marcia Pereira de Oliveira Duarte, Luciana Polaco Covre, Daniel Claudio de Oliveira Gomes, Alda Maria da Cruz, Alessandra Marcia da Fonseca Martins, Herbert Leonel de Matos Guedes

**Author notes:** Corresponding Author: Dr. Herbert Leonel de Matos Guedes, Universidade Federal do Rio de Janeiro, Instituto de Microbiologia Paulo de Góes, Grupo de Imunologia e Vacinologia, Laboratório de Imunobiotecnologia, Av. Carlos Chagas Filho, 373, CCS, Room I2-043, Rio de Janeiro, 21941-902. RJ, Brazil., Laboratório de Imunologia Clínica, Instituto Oswaldo Cruz, Fundação Oswaldo Cruz, Rio de Janeiro, RJ, Brazil; Postal Code: 21040-360.

## Abstract

Mucosal Leishmaniasis is one of the most aggressive clinical manifestations of *Leishmania* infection disease, characterized by the destruction of nasal and oral tissues. The mechanisms by which this disease occurs are still not well understood due to the lack of effective experimental models. Mucosal leishmaniasis is associated with inflammatory response, especially Th17 response. Based on that, in this work, the immunopathological aspects of the experimental infection of BALB/c mice by *Leishmania amazonensis* in the mucosa site were evaluated due to high susceptibility and the disease been associated with Th17 response. Three infection modes were performed and compared according to the injection site. Six weeks post infection, mice presented edema in the nasal and premaxillary region, with progressive growth until twelve weeks. The micro–Computerized Tomography and the histology images demonstrated that the parasite inoculation led to destruction of squamous and transitional tissues in NC and NB groups, with several cells harboring amastigotes. Mice infected in the mucosa tissues had higher parasite load and IgG, IgM antibody levels and increased production of cytotoxic mediators such as CD107, granzyme b and perforin, inflammatory cytokines as IFN-γ and IL-17, but lower frequencies of CD4^+^ IL-10^+^ cells compared to ear dermis. Taken together, our data shows that *L. amazonensis* parasites are more proliferative in nasal mucosa and the infection leads to an increased inflammatory response compared to ear dermis, pointing to this model as an interesting approach to understand some features of MCL immunopathology.

## Introduction

Leishmaniasis is a group of neglected tropical diseases caused by infection by protozoan parasites belonging to the genus *Leishmania*. The disease affects the most deprived populations in the world, being endemic in more than 98 countries, with cases distributed across Asia, Africa, the Middle East, and South and Central America [1]. Estimates suggest that around 1 million new cases of leishmaniasis occur every year globally, with a population of 1 billion people living in areas at risk of contracting the disease [2].Of the different clinical manifestations of leishmaniasis, mucocutaneous leishmaniasis (MCL) stands out because it can generate a severe deformity, generating destruction of the tissues of the nose, upper lips, palate, pharynx, larynx, perforation of the septum and loss of bone tissue [3, 4]. Mucocutaneous Leishmaniasis (MCL) is a consequence of *Leishmania*’s tropism for mucosal tissues, involving the respiratory mucosa of the upper tract and the oral cavity. This clinical manifestation is typically a result of infection with New World species such as *Leishmania braziliensis*, *Leishmania panamensis*, *Leishmania guyanensis* and *Leishmania amazonensis* [5]. It is estimated that 3 to 5% of cases of cutaneous leishmaniasis caused by these species evolve into the mucosal form [6]. The cure rate for MCL treatment is lower compared to LCL [7, 8], which highlights the importance of a better understanding of development and immune response in this clinical manifestation [9, 10].

Therefore, despite being a more severe manifestation and even with a wide range of *Leishmania* species that can cause MCL and capable of infecting small rodents such as hamsters and mice, there is a lack of experimental models for MCL [5], and the immune response associated with this clinical manifestation is still poorly characterized. Using *L. amazonensis* infection on BALB/c mice that is very susceptible to infection, we hypothesized that the parasite could establish the infection directly to the nose inducing a manifestation similar to mucosal leishmaniasis. Previous attempts were made in the literature using subcutaneous infection in the paw leading to mucosal metastasis, however, they are very long (approximately 8 months after infection) diverse (some animals do not develop the disease) and several mice don’t survive during the prolonged infection and aging process. It is urgent a model that facilitates the study of mucosal leishmaniasis [11]. In the context of the disease, mucosal leishmaniasis is associated with a Th1, but also a Th17, neutrophil infiltrate immune response [12]. Meanwhile, BALB/c dermal pathology is associated with Th2 [13] but also with a strong Th17 [14, 15], making BALB/c a model to exploit the role of those immune axis to the pathology of mucosal leishmaniasis. In this way, here in we established a *L. amazonensis* mucosal leishmaniasis mice model using three infections sites to understand the immunopathology of the disease.

## Methods

### Leishmania Culture

In this work, parasites of the species *L. amazonensis* (MHOM/BR/75/JOSEFA) were used. The promastigotes were maintained in 25cm² culture flasks with M199 medium (pH 7.2), supplemented with 10% fetal bovine serum, 5µg/mL bovine hemin, 50U/mL penicillin, and 50µg/mL streptomycin.

### Animals

Mice from BALB/c lineage, females aged 8-12 weeks, were used. The mice came from the Biotério Central de Camundongos of the Centro de Ciências da Saúde (CCS) from the Universidade Federal do Rio de Janeiro, mice were maintained under the Protocol of the Ethics Committee on the Use of Experimental Animals No. 133/23 of the CCS.

### Infection and Lesion Development

The parasites were cultivated until the beginning of the stationary phase. The culture containing the promastigote forms were collected, washed with phosphate buffer saline (PBS); and centrifuged for 10 minutes at 4°C at 1000 x G force. After centrifugation, the supernatant was discarded, and the cells were washed with PBS. The procedure was repeated twice. Then the cells were counted and adjusted to a concentration of 2×10^8^ parasites/mL. The animals were sedated with ketamine and xylazine and infected with 2×10^6^ parasites using Hamilton® syringes coupled with 34G needles. Different infection methods were implemented: intradermal in the ear performed at an angle of 20°; Cutaneous Nose, with the inoculum carried out at the apex of the mouse’s nose; Septum, with the inoculum occurring in the mouse septum and “Nasobasal”, with the inoculum being carried out at an angle of 60° towards the lower “floor” part of the nose. The lesion was monitored with photos. Weekly measurements of the mice’s ears were taken using Mitutoyo™ thickness gauges.

### Micro Computerized Tomography, 3D Reconstruction e Lesion Analysis

Animals were sedated and placed in the LabPET8 Flex Triumph Gamma Medica™ tomography system at CENABIO, UIPA. The equipment settings included 60kV, 480µA, 1024 projections, 8-minute acquisition, and 30x magnification with a 39.46mm field of view, focusing on the snout and mucous membrane. After acquisition, data was reconstructed and analyzed using 3D Slicer™ software, with the Scissors tool applied to generate the snout area.

### Histology Analysis

Tissues from animal snouts were fixed in 4% formaldehyde in PBS for 15 days. Decalcification was performed for 28 days using 10% EDTA. After fixation and decalcification, tissues were dehydrated through a series of ethanol and xylene solutions, then embedded in paraffin. Sections (5µm thick) were cut, rehydrated, and stained with Hematoxylin & Eosin (H&E). Tissue processing was done by the Histotechnology Platform at IOC - FIOCRUZ. Histology was performed with n = 5 per group.

### Parasite Load Quantification

At the end of each experiment, parasite load was assessed using the limiting dilution method (LDA). Animals were euthanized with an overdose of Ketamine and Xylazine, and tissues (snout, nose tip, cheek, ear, spleen, NALT, auricular and cervical lymph nodes) were collected, weighed, and macerated in 1mL of M199. Serial 1:4 dilutions were made, and the results were plotted. Three experiments (n=3-5) were conducted, with two examining the entire snout and one focusing on segmented parts, 12 weeks post-infection.

### Flow Cytometry

Flow cytometric analysis of cells from cervical and ear lymph nodes of infected or naïve animals was performed. Tissues were macerated, and cells counted with 0.1% trypan blue dye. 1×10^6^ cells were plated in a 96-well plate, centrifuged, and stimulated with PMA (20ng/mL), ionomycin (1µg/mL), and monensin (2.5µg/mL) in Complete RPMI for 4 hours at 37°C. Cells were blocked with anti-FcR, surface stained, fixed using the eBioscience™ FoxP3 kit, and intracellularly labeled. The samples were analyzed on a BD LSR Fortessa X-20 cytometer. Table S1 denotes the cytometer configuration. Table S2 shows the antibodies (BioLegend™) used and their concentrations. Figure S1 denotes the performed gating strategy.

### Statistics

Results are represented by Standard Error of the Mean (SEM). Statistical significance was performed by unpaired two tailed Student t test with 95% confidence interval (p<0,05) for the respective data: Lesion profile by tomography, parasite load quantification and cytometry. While One Way ANOVA with 95% interval of confidence (p<0,05) was performed using Tukey post-test, for the ELISA OD Sum data. Statistically significant differences were defined as * for p<0,05; ** for p<0,005; *** for p<0,0005.

## Results

### Direct inoculation of *Leishmania amazonensis* in nasal mucosa led to injury but the lesion profile depends on the type of inoculum performed

To assess *Leishmania* infection in the nasal mucosa, 2×10^6^ promastigotes of *L. amazonensis* were directly inoculated into different nasal sites of mice. After 6 weeks, the Nose Cutaneous (NC) and Nasobasal (NB) groups developed visible lesions with edema and redness (figure 1A). These lesions progressed and became necrotic by 12 weeks in some cases. The muzzle volume was significantly larger in the NC and NB groups (387mm³ and 616mm³) compared to the Sham group (figure 1B), resembling the progressive ear infection, which reached 1.8mm in thickness (figure 1C, 1D). In contrast, the Septum group showed no visible lesions throughout the study (figure 1A). These results indicate that nasal inoculation can cause tissue damage, depending on the infection site.

**Figure 1.**
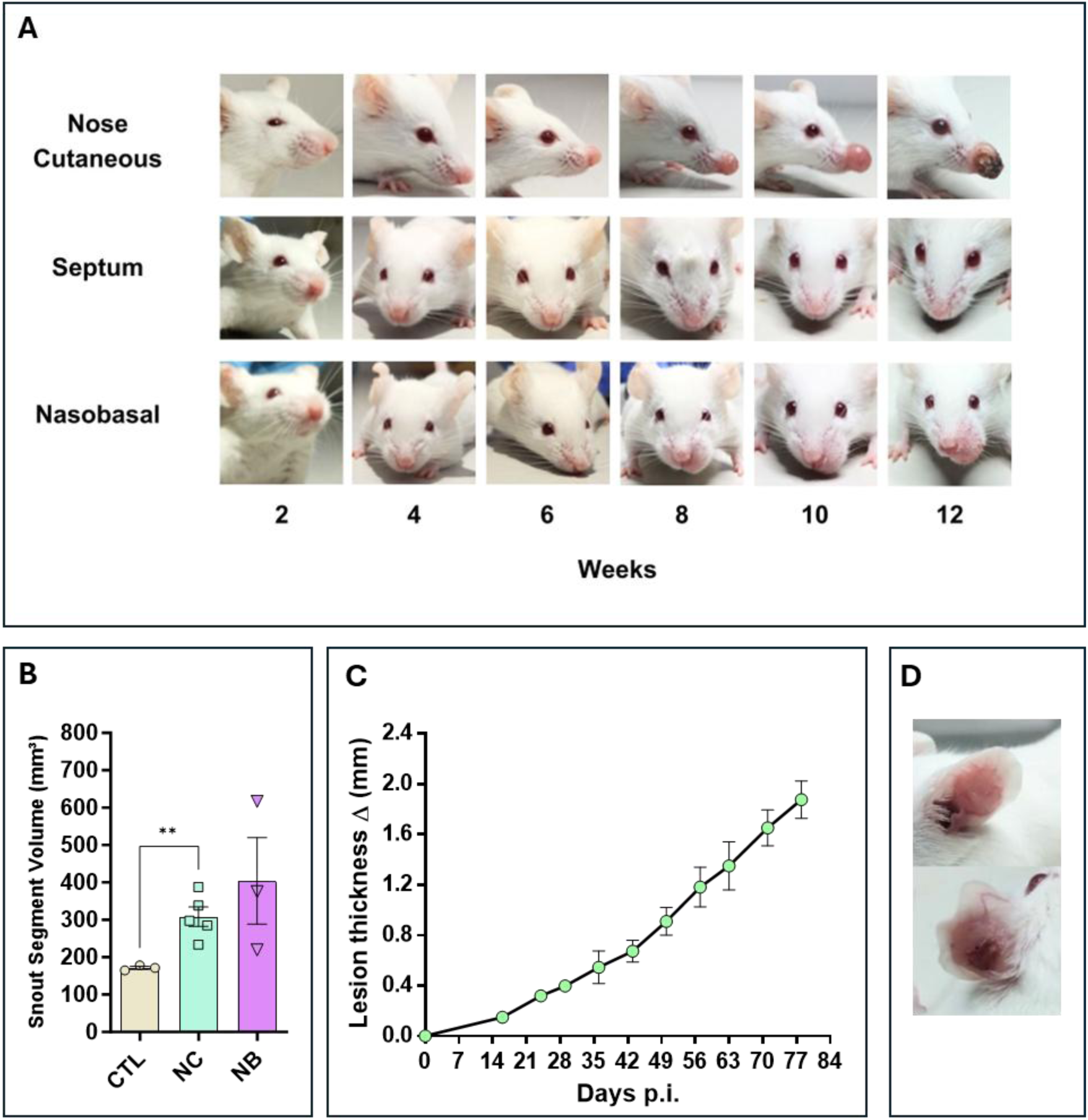
Nasal mucosa lesion profile. The modes of nasal mucosa inoculation with *L. amazonensis*. Nose cutaneous (NC) group was inoculated in the nostril nose dermis; Septum (ST) was inoculated in the septum tissue, “Nasobasal” (NB) was inoculated in the nasal floor. In all cases was used an 34G needle. **(A)** Lesion profile of the three groups, during a period of 12 weeks post infection (p.i), showing a continuous development of edema and erythema starting at 6 to 8 weeks for the NC and NB group. ST group does not show any sign of lesion. **(B)** Snout volume quantification 12 weeks p.i (Standard Error of the Mean – (SEM)), using micro–Computerized Tomography (micro-CT). NC and NB groups have greater volume compared to control groups. **(C)** Ear dermis lesion thickness of mice infected in the ear as an infection control group (SEM). The thickness continuous increases during the experiment reaching around 2.0mm in 12 weeks p.i. **(D)** Representative ear of a mice from control infection group showing edema and erythema. (B) Representative data of 3 independent experiments (5 animals per group). (C) Data from one experiment 3-5 animals per group. (D) Representative data from 2 independent experiments 5 animals were used. Statistics: unpaired t-test * p < 0.05.

### The Nasal mucosa model showed a higher parasite load than ear dermis infection

Parasite load in infected mice ranged from 10^6^ to 10^8^ parasites per gram of tissue in the Nose Cutaneous (NC) and Nasobasal (NB) groups, while the Septum (ST) group showed only 10^4^ parasites, with Only 20% of animals testing positive (Figure 2A). Ear dermis infections had 10^6^ parasites per gram (Figure 2B). Parasites were mostly concentrated in the edema sites, such as the nose tip and pre-maxillary regions (Figures 2C, 2D). Some animals in the NB group had around 10^5^ parasites in NALT (Figure 2E). Only one animal in the NC group showed spleen infection (Figure 2F). The cervical lymph nodes of NC and NB mice had 10^6^ parasites (Figure 2G), while the ear lymph nodes had 10^5^ parasites (Figure 2H). The Septum group showed parasitic load in the cervical lymph node but not the nasal cavity (Figure 2G). These results indicate that mucocutaneous *L. amazonensis* infection leads to a higher parasite content in nasal mucosa infection compared to ear dermis, with the cervical lymph node being the primary draining node.

**Figure 2.**
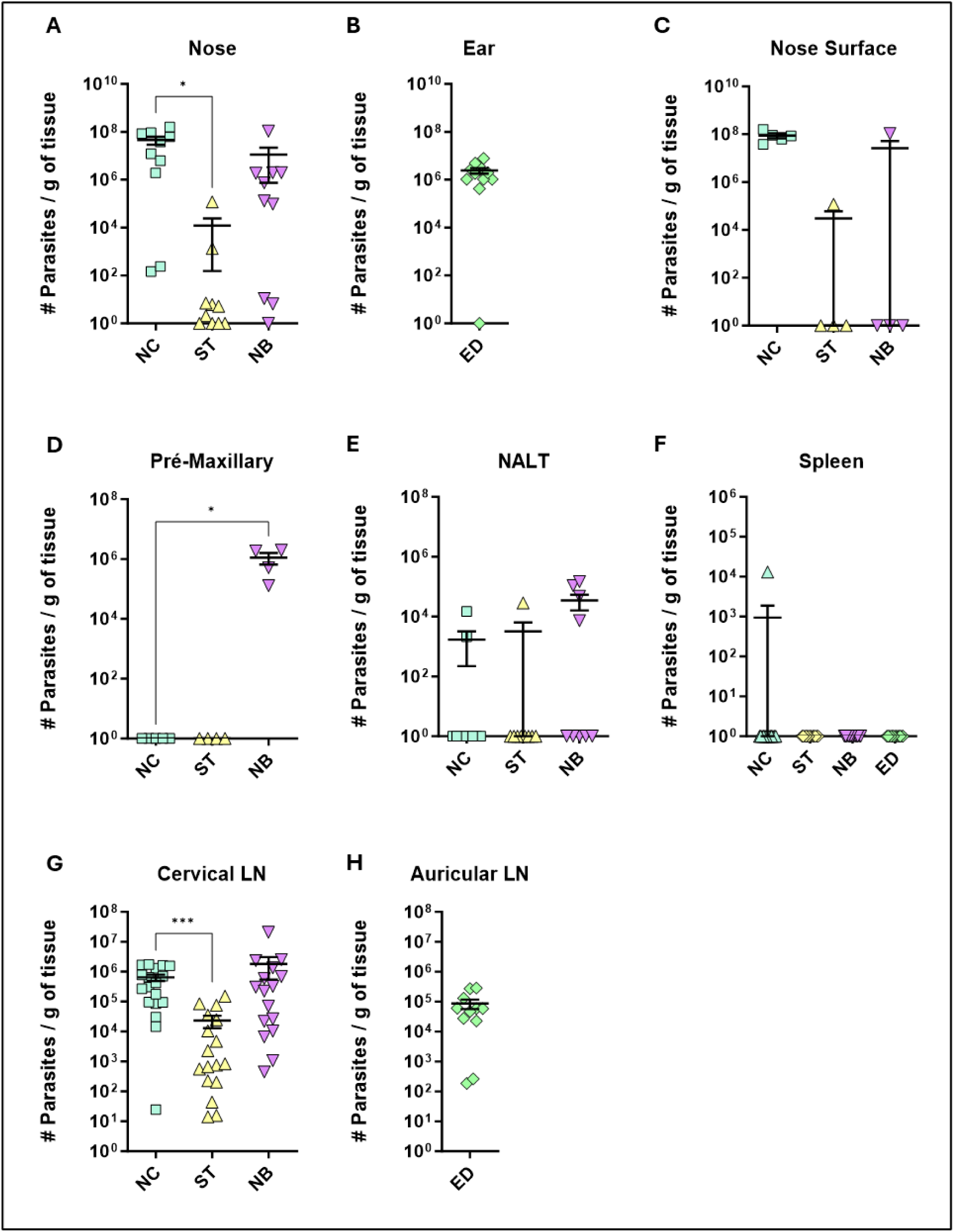
Parasite Load Profile. Profile of parasite load per gram of tissue, of the different groups of infection. NC, ST, NB, ED means respectively to Nose Cutaneous, Septum, “Nasobasal” and Ear dermis infection groups. The following tissues are represented in the figure: **(A)** Nose = the entire snout containing the nasal tissues; **(B)** Ear; **(C)** Nose Surface = only the visible nose; **(D)** mice premaxillary region; **(E)** Nasal Associated Lymphoid Tissue (NALT); **(F)** Spleen; **(G)** Cervical lymph node; **(H)** auricular lymph node. Parasite load was detected by the limiting dilution method (LDA) carried out at the end of the experiment (approximately 12 weeks of infection). (A) Data from 3 independent experiments; (B, C, and D) 1 independent experiment; and (E), Data from 4 independent experiments; (G and H), 2 independent experiments. N = 5 animals per group. Data represented by the Standard Error of The Mean (SEM). Statistics: unpaired t-test * p < 0.05.

### Nasal infection induced nasal swelling, but no septum perforation or new nasal cavity formation took place

MicroCT analysis was performed to assess nasal cavity impairment, focusing on the NC and NB groups, as the ST group showed no visible lesions or significant parasite load. NC-infected mice typically had lesions at the nose apex, while NB-infected mice had lesions affecting the mucosa and premaxilla, extending to the upper lips (Figure 3A, 3B). No nasal cavity enlargement, perforations, or deformation were observed compared to controls (Figure S2), but tissue swelling was evident in both NC and NB groups (Figures 3A-D). Axial sections showed noticeable edema on the outer nasal cavity (Figure 3C, 3D). These results indicate that lesion profiles depend on the inoculation site, mainly affecting the nasal tissues initial portions.

**Figure 3.**
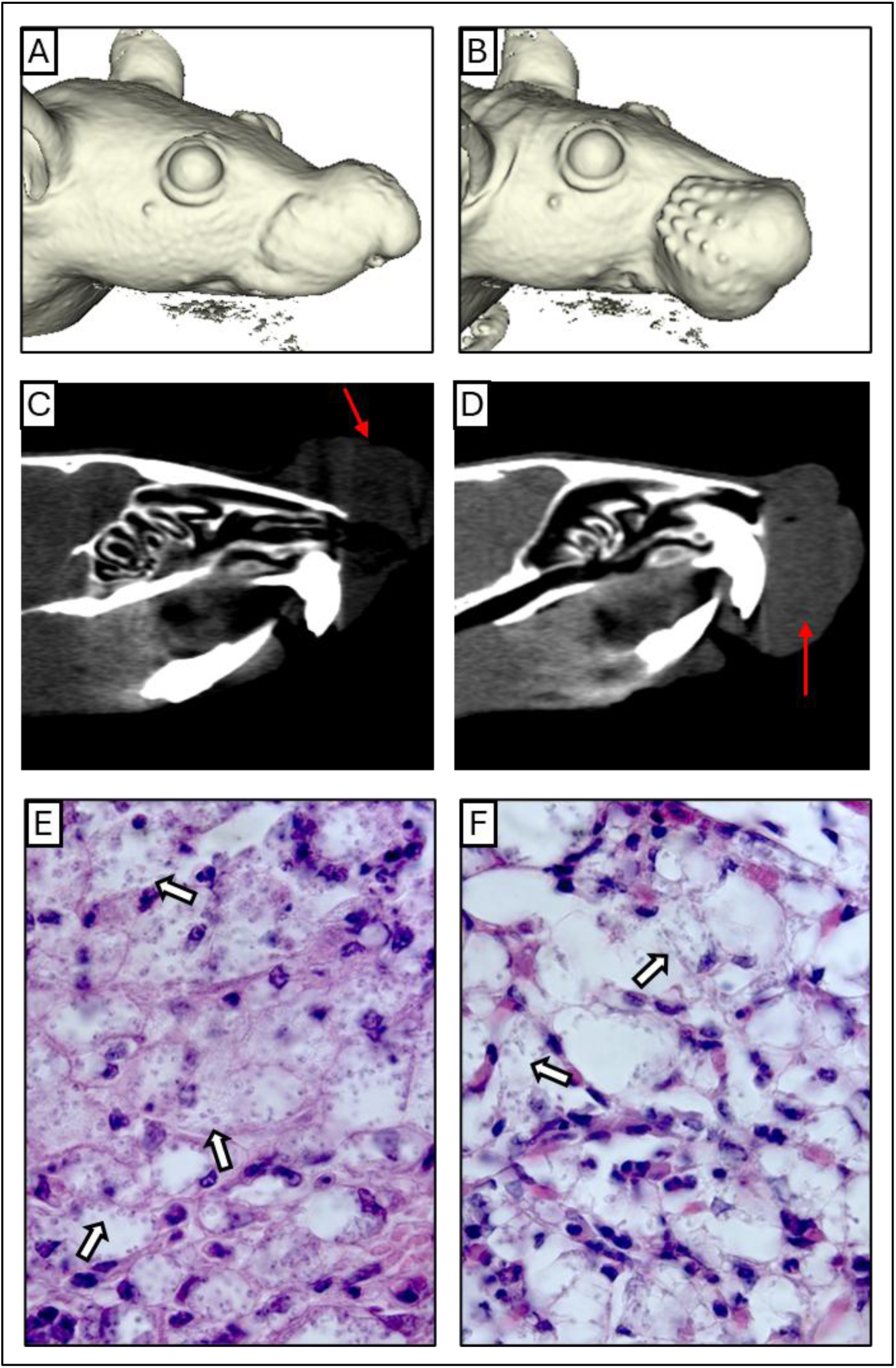
Anatomical and Histological Profile. **(A,B)** A complete three-dimensional reconstruction of NC and NB mice model respectively. **(C, D)** Coronal Section planes showing edema and partial nose obstruction of NC and NB mice respectively. **(E, F)** Intense inflammatory infiltrate containing infected cells, below the epithelium of NC and NB mice respectively. The red arrows indicate the region of edema, and the white arrows show examples of cells bearing amastigotes. Magnification: E) 1000x and (F) 1000x.

### Nasal infection promoted destruction and deconfiguration of squamous and transitional tissues

Histological analysis of the nasal cavity revealed significant inflammation and parasite presence. In the NC group, edema with a neutrophilic infiltrate and vacuolated macrophages containing amastigotes was observed (Figure 3E, S3), along with respiratory epithelium loss (Figure S4B), and parasitized macrophages near cartilage (Figure S4E). In the NB group, similar edema and neutrophil infiltration were seen, with many parasitized macrophages (Figure 3F). Respiratory epithelium showed slight morphological changes, including cellular stress and detachment (Figure S4C, S4F). These findings indicate that *Leishmania* infection impacts the inner nasal mucosa in this model.

### Nasal mucosa infection produced higher systemic antibodies titters compared to ear dermis infection

To assess immune responses to different nasal mucosa infections by *Leishmania*, we measured antibody levels. The NC group showed the highest IgM, IgG, and IgG1 levels, followed by NB and ED (Figure 4A-F). IgG2a and IgG2b were highest in NB, then NC (Figure 4G-J). Overall, NC and NB had stronger antibody responses than ED, while ST had the lowest. These findings suggest mucocutaneous infection triggers a stronger humoral response than ear dermis infection, with antibody profiles varying by inoculation site.

**Figure 4:**
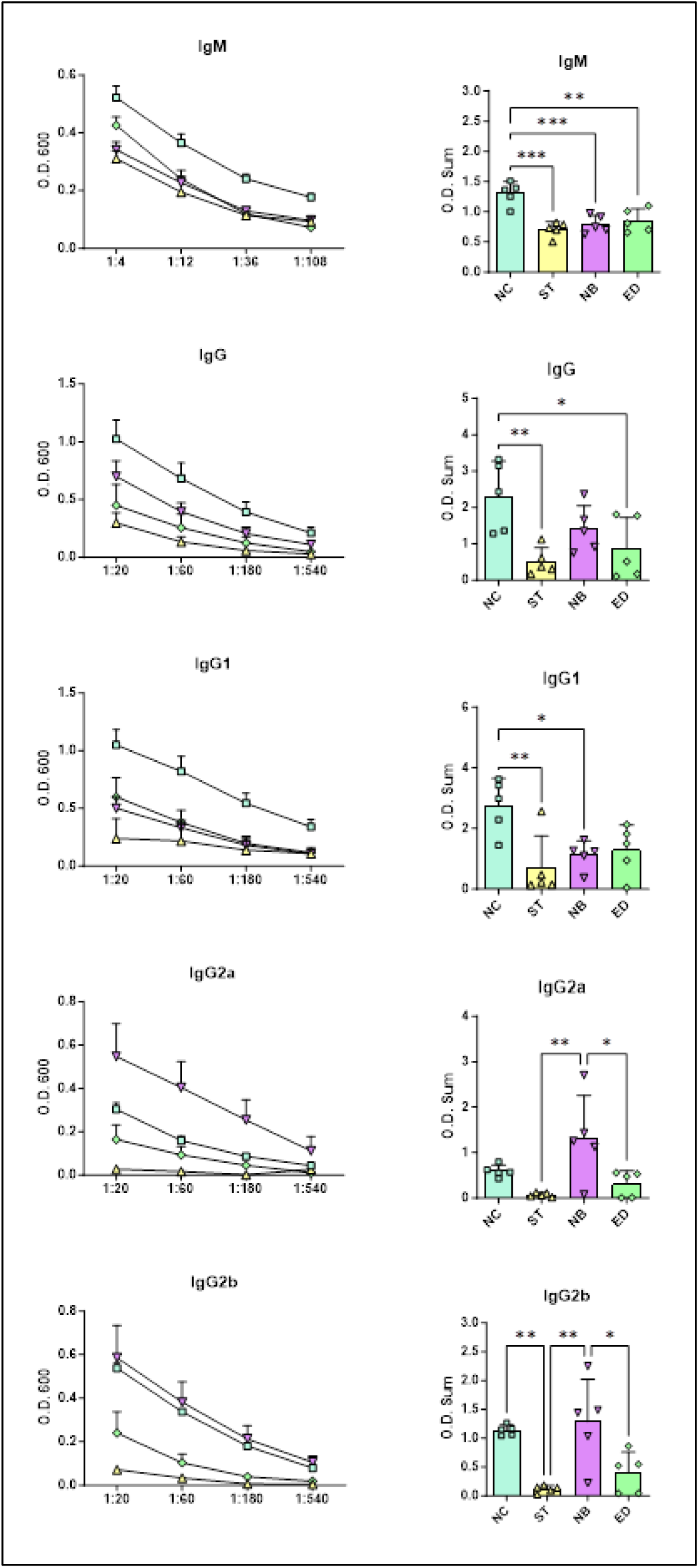
Systemic antibodies infection profile. Systemic antibodies responses in the different infected groups. **(A, C, E, G)** Ig titration using four dilutions, represented by O.D. 600 absorbance measurements (Standard Error of The Mean (SEM)). **(B, D, F, H)** Ig O.D. Sum from each animal and groups. NC groups produce more IgM, total IgG and IgG1, while NB mode of infection have more IgG2a and IgG2b. NC, ST, NB, ED groups represented by square; triangle; inverse triangle and diamond respectively. Statistics: One-way ANOVA; * p<0,05; ** p<0,005; ***p<0,0005.

### Cytotoxic and effector CD4^+^ and CD8^+^ T cells from draining lymph nodes were increased in NC, however, CD4^+^ T cells producing IL-10 were impaired in nasal mucosa infection

Flow cytometry of draining lymph nodes showed reduced CD4+ T cell frequency in the ED group but not in nasal infections. The NC and NB groups had increased total CD4+ T cells (Figure 5A) and higher numbers of cells expressing cytotoxic markers (CD107a, granzyme B, perforin) across NC, NB, and ED groups (Figures 5B-D), despite unchanged frequencies (Figures S6-S8).

**Figure 5.**
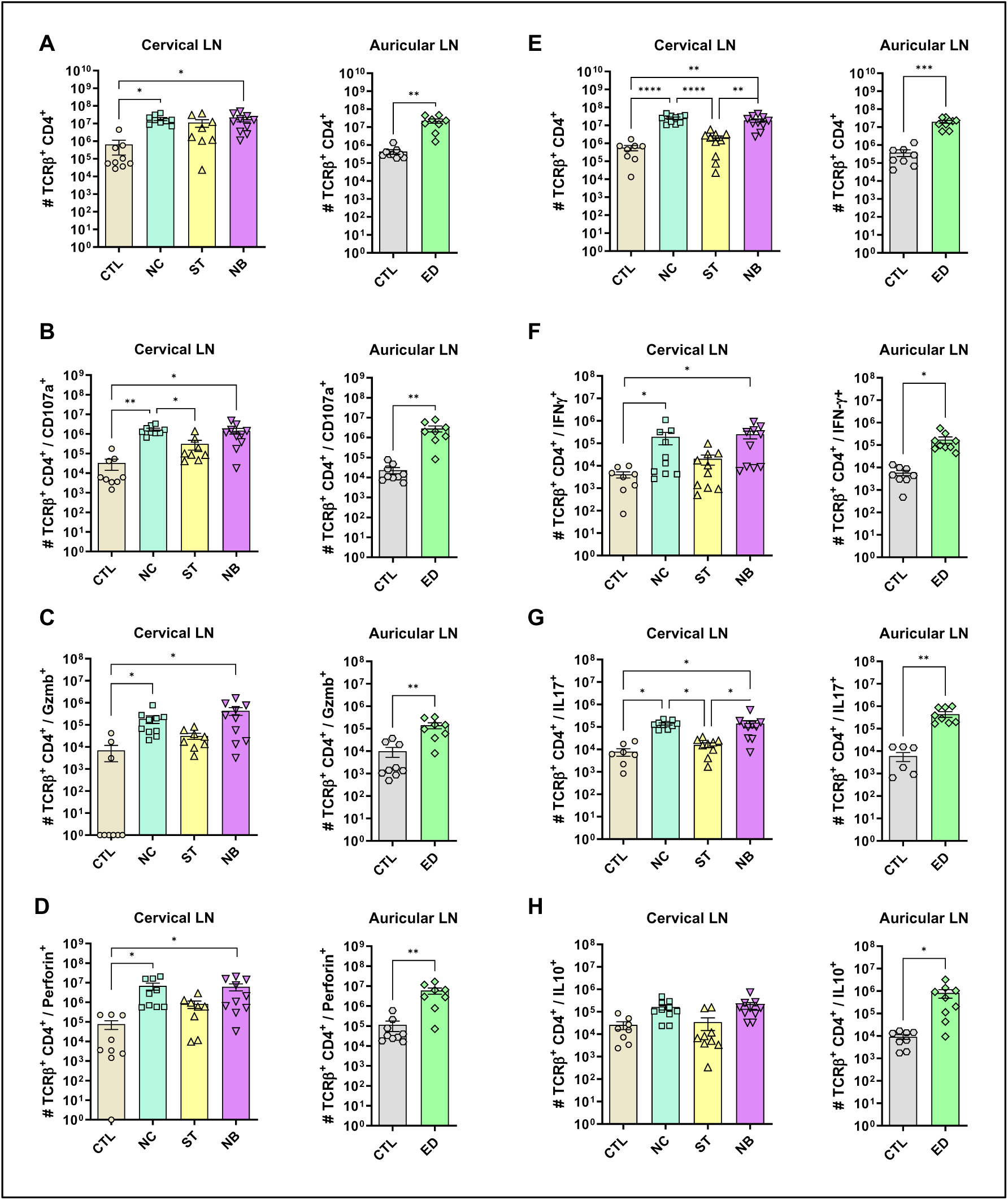
CD4^+^ T cells cytotoxic and effector profile. (A-D) The cytotoxic profile of T helper lymphocytes was measured by the expression of CD107a, Granzyme B and Perforin. (A) Cervical and auricular lymph nodes total Lymphocytes counts are shown in logarithmic scale; (B) total cells expressing CD107a, (C) Granzyme B; (D) Perforin. (E-H) The effector profile of T helper lymphocytes was measured by the expression of IL-10, IL-17 and IFN-γ. (E) Cervical and auricular lymph nodes total lymphocytes counts are shown in logarithmic scale; (F) total cells expressing IFN-γ, (G) IL-17; (H) IL-10. Data accumulative of two independent experiments (3-6 animals per group). Statistics: plot with Standard Error of The Mean (SEM), t-test was used for all groups and samples * p<0,05, **<0,005.

Although frequencies remained unchanged (Figures S9-S10), total IFN-γ+ (Figure 5F) and IL-17+ (Figure 5G) cells increased in NC, NB, and ED groups. CD4+ IL-10+ T cells increased only in the ED group (Figure 5H), with lower IL-10-producing CD4+ T cell frequencies in nasal infections compared to ED (Figure 6A, B), highlighting distinct expansion dynamics between nasal and dermal infections.

**Figure 6.**
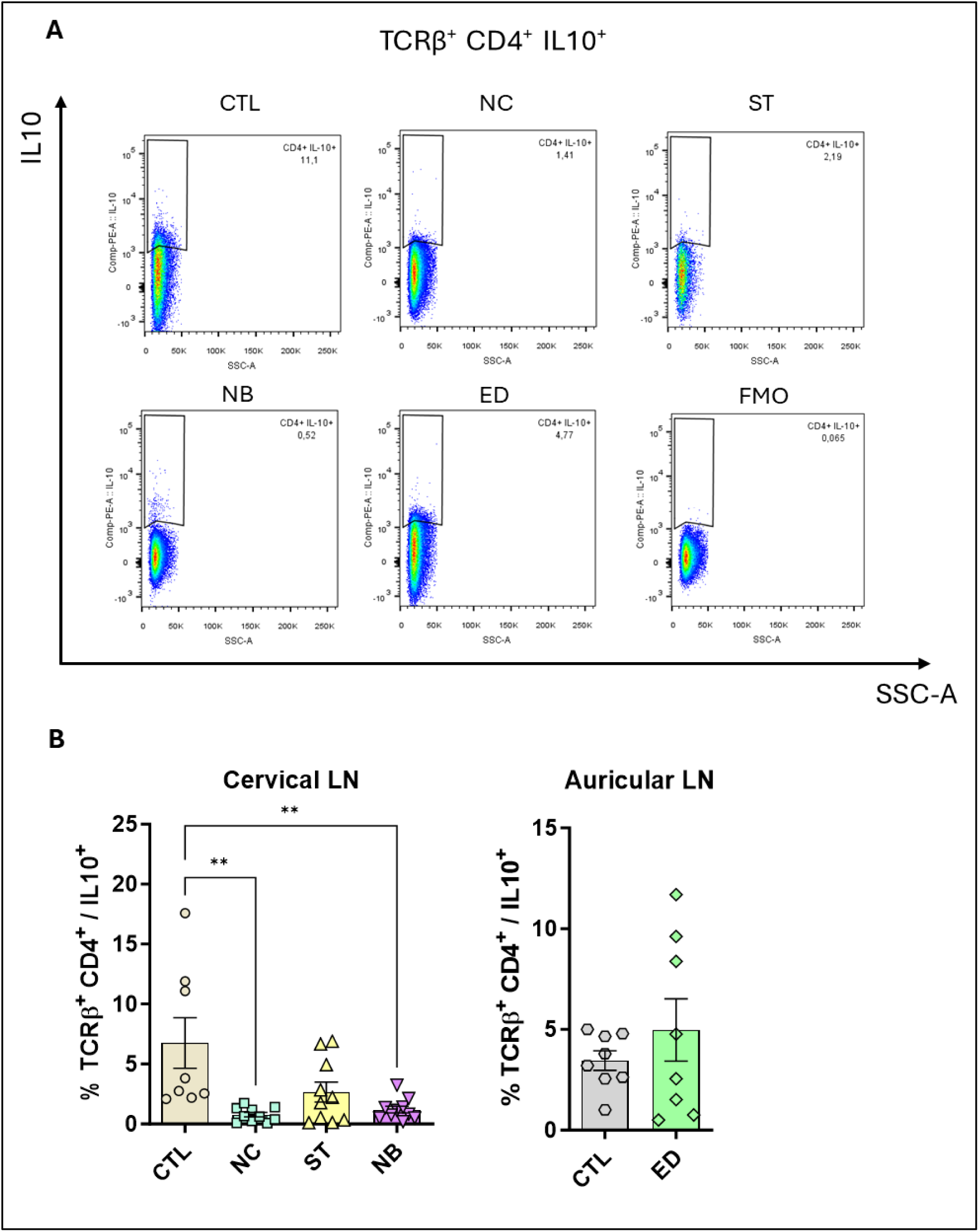
CD4^+^ T cells IL-10^+^ is impaired. The IL-10 producing CD4^+^ T cells was measured, and the frequency is shown. (A) Pseudocolor dot plot showing the representative animals of each group. (B) Frequency plot from cervical and auricular lymph nodes from infected animal vs controls. Data accumulative of two independent experiments (3-6 animals per group). Statistics: plot with Standard Error of The Mean (SEM), t-test was used for all groups and samples * p<0,05, **<0,005.

We analyzed whether changes in CD4+ IL-10+ T cells were linked to CD4+ CD25+ FoxP3+ Tregs. Treg numbers increased in NC, NB, and ED groups without frequency changes (Figure S11). Similar trends were observed for CD25-FoxP3+, CD25+ FoxP3-, and CD25-FoxP3-cells (Figures S12-14), indicating the IL-10 reduction is unrelated to regulatory T cell frequency.

We also analyzed CD8^+^ T lymphocytes during mucosal infections. NC and NB groups showed an expansion in total CD8^+^ T cells (Figure 7A) with frequencies similar to CD4^+^ T cells (Figure S15). Cytotoxic CD8^+^ T cells markers increased in NC, NB, and ED groups (Figures 7B-D), with NC showing higher CD8^+^ Perforin^+^ frequency (Figure S18), nut no frequency changes to CD107a and Granzyme B respectively (figure S16, S17). There were no IFN-γ^+^ CD8^+^ significant changes in frequency (Figure 7F) or number (Figure S19). IL-17^+^ CD8^+^ T cells increased in NB and ED groups (Figure 7F) without frequency changes (Figures S20). CD8^+^ IL-10^+^ T cells showed no numerical change (Figure 7H), but frequency decreased in mucosa-infected and ED groups (Figure S21).

**Figure 7.**
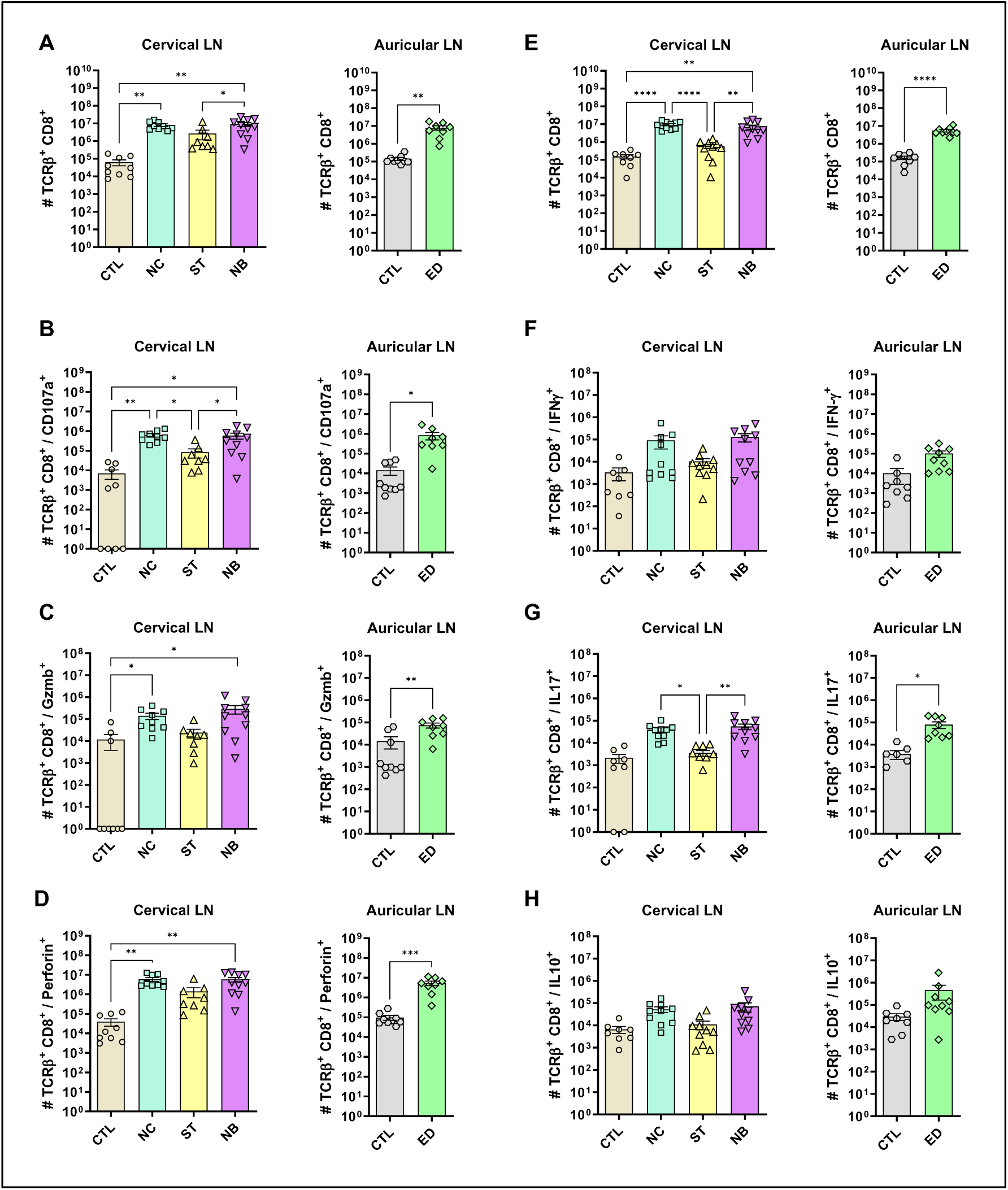
Cytotoxic and effector profile of CD8^+^ T cells. (A-D) The cytotoxic profile of T helper lymphocytes was measured by the expression of CD107a, Granzyme B and Perforin. (A) Cervical and auricular lymph nodes total Lymphocytes counts are shown in logarithmic scale; (B) total cells expressing CD107a, (C) Granzyme B; (D) Perforin. (E-H) The effector profile of T helper lymphocytes was measured by the expression of IL-10, IL-17 and IFN-γ. (E) Cervical and auricular lymph nodes total lymphocytes counts are shown in logarithmic scale; (F) total cells expressing IFN-γ, (G) IL-17; (H) IL-10. Data accumulative of two independent experiments (3-6 animals per group). Statistics: plot with Standard Error of The Mean (SEM), t-test was used for all groups and samples * p<0,05, **<0,005, ***<0,0005.

## Discussion

Mucocutaneous Leishmaniasis (MCL) causes severe tissue damage in nasal, oral, and pharyngeal mucosa, leading to edema and erythema in affected areas [3, 5]. In our mouse model, the NC and NB groups developed visible edema and erythema by week 6, progressing over time. Lesions in the NC group formed at the nose tip and extended dorsally, resembling the experimental mucosal leishmaniasis in dogs and hamsters [16, 17]. In the NB group, lesions spread to the upper lips and premaxilla, mimicking clinical features of human MCL [18, 19].

Some animals developed necrotic lesions after 12 weeks of infection. While *L. amazonensis* typically causes non-necrotizing lesions in BALB/c mice resembling diffuse cutaneous leishmaniasis in humans [20]. Similar variation in infection sites was observed with *Leishmania major* [21], possibly linked to site-specific microbiota. Nasal microbiota, including *Staphylococcus* and *Streptococcus* species [22, 23], may exacerbate tissue damage by recruiting neutrophils and CD8 IL-17+ cells during infection [24, 25].

Mucosal leishmaniasis in humans often affects septa, cartilage, and nasal cavities, with occasional bone destruction visible via computed tomography [3, 26–28]. In this study, microCT revealed edema in the nasal and premaxilla regions of NC and NB groups. Histopathology showed dense cellular infiltrates and infected cells near cartilage and the nasal mucosa, with respiratory epithelium changes, consistent with previous findings [11].

Despite MCL caused by *L. braziliensis* typically shows fewer parasites at the site than cutaneous leishmaniasis [29], our model NC and NB groups had higher parasite loads, including in draining lymph nodes and infected macrophages in nasal mucosa. This may reflect *L. amazonensis* characteristics, which produce higher parasite loads than *L. braziliensis* in mice [30] as evidenced also by the the cases o MCL caused by *L. amazonensis*, where parasite amastigotes were described as easily visualized in H&E sections [31]. Unlike previous models focusing on nasal infection [16, 17, 32, 33], this study uniquely examined infections within the nasal mucosa cavity, revealing that the inoculum site significantly influences lesion severity and parasite load across NC, ST, and NB groups. Necrosis was observed in some animals, a feature absents in other models [16, 17, 33]. Mucosal infections showed clinical aspects similar, or more severe than cutaneous ear lesions, aligning with findings in *L. panamensis*-infected hamsters [17]. While other models required approximately 8 months to develop lesions [11], this approach shortened progression to 6–8 weeks. Future studies could compare axenic amastigote infections, given their higher virulence and differing proliferation kinetics from promastigotes [34, 35].

MCL severity correlates with higher anti-*Leishmania* IgG levels [36]. In our model, mucosal infections (NC and NB) produced more IgM and IgGs than dermal infections, with NC favoring IgG1 (Th2 response) and NB favoring IgG2a (Th1 response). This highlights site-dependent immune response differences, consistent with established antibody production dichotomy [37], and the high antibody levels produced by the mucosal infection compared to dermis can be interpreted as an increased inflammatory response that may contribute to the severity of pathology.

To compare nasal mucosa infections with human MCL, we performed flow cytometry on lymph node cells. Mucosal infection (NC and NB) increased CD4+ and CD8+ T cells expressing cytotoxic markers (CD107a, Granzyme B, Perforin) and IL-17, with less IFN-γ, but no increase in IL-10-expressing cells, unlike the ED group. In MCL, CD4+ T cells produce high IFN-γ and TNF-α, with elevated Th17 responses that recruit neutrophils and cause tissue damage [12, 38, 39]. MCL also shows higher granzyme A-expressing cells, linked to greater damage [34], with a lower IL-10 receptor expression, despite normal IL-10 expression [39]. Our data suggests a similar imbalance between cytotoxic and regulatory responses, with reduced IL-10+ cells and increased IL-17+ cells potentially contributing to severity, as IL-10 limits Th17-mediated pathology [40].

Taken together, *L. amazonensis* infection in BALB/c mice induces an immune response that reflects some aspects of the human MCL, with increased cytotoxic and effector cells but a compromised regulatory response. However, our studies didn’t show some MCL features like cavity formation, septal perforations, and bone destruction, possibly due to parasite species or host lineage differences, or the shortened pathology compared to human pathology [28]. To address this, ongoing research is being performed with C57BL/6 strain for a stronger Th1 response [15, 41]. We also are planning to test the *L. braziliensis* hamster model for its higher susceptibility for this species [42]. As NC and NB sites differed in some characteristics, we believe that the combination of those inoculum sites may better represent the full mucosal leishmaniasis profile in mice. More studies are needed to confirm the lymphocyte profile in the mucosal site.

In conclusion, when infected directly at mucosal site, *L. amazonensis* parasites are more proliferative in nasal mucosa and lead to an increased cytotoxic and antibody response compared to ear dermis infection, which can be associated with the IL-10 impairment in the nasal mucosa that can be related to the immunopathology of MCL.

## Author Contribution

Investigation: Alisson Amaral da Rocha^1,2^, Júlio Souza dos Santos^1,2^; Alessandra Márcia da Fonseca Martins^1,2^; Igor Bittencourt dos Santos^1,2^; Douglas Barroso de Almeida^1,2^; João Victor Paiva Romano^1,2^; Naiara Carla dos Santos Manhães^1,2^; Hozany Praxedes dos Santos^1,2^; Marcia Pereira de Oliveira Duarte^4^; Antônio José da Silva Gonçalves^4^.

Formal Analysis: Alisson Amaral da Rocha^1,2^; Alessandra Márcia da Fonseca-Martins^1,2^ Elias Barbosa^3^

Conceptualization and Methodology: Alisson Amaral da Rocha^1,2^, Júlio Souza dos Santos^1,2^ Herbert Leonel de Matos Guedes^1,2^

Resources: Herbert Leonel de Matos Guedes^1,2^, Alessandra Márcia da Fonseca-Martins^1,2^, Alda Maria da Cruz^4^, Daniel Claudio de Oliveira Gomes^5^, Luciana Polaco Covre^5^,

Writing – Original Draft: Alisson Amaral da Rocha^1,2^

Supervision and Funding Acquisition: Herbert Leonel de Matos Guedes^1,2^

## Acknowledgement

We would like to acknowledge the Centro Nacional de Bioimagem (CENABIO) for the equipment and technical support in tomography analysis. We would also like to acknowledge the Platform of Histotechnology (FIOCRUZ-RJ) for the processing of histology samples.

## Supplementary Material

**Figure S1.**
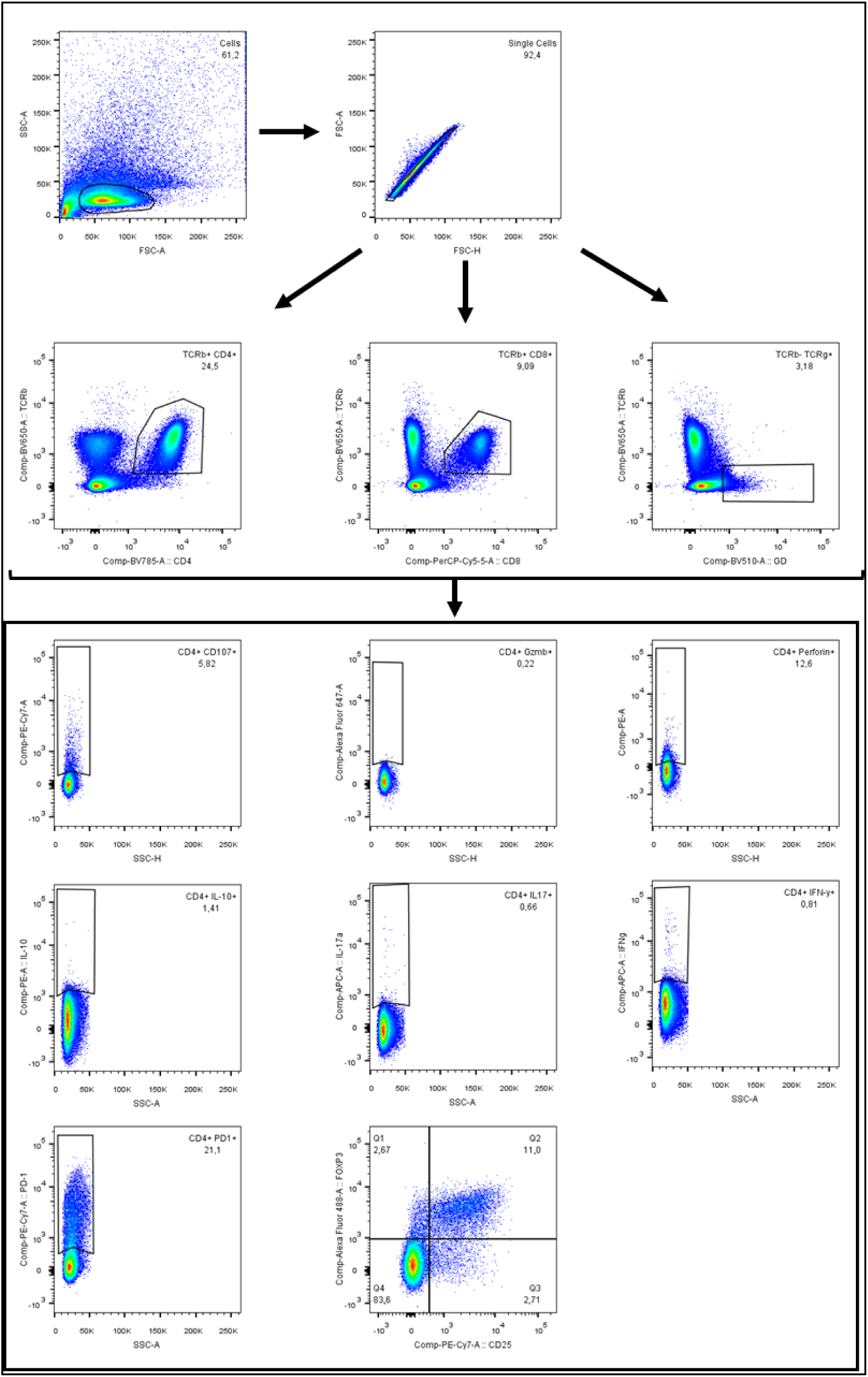
Gating strategy. The figure shows the gating strategy used to find and analyze CD4^+^ T, CD8^+^ T and Tγδ^+^ T cells. Within each population, a gate was performed to highlight cells expressing the molecules CD107a, Perforin, Granzyme B, IL10, IL17 and IFN-γ (except Tγδ^+^) and PD-1. In CD4^+^ T cells, the concomitant expression of CD25 and FoxP3 was also verified.

**Figure S2.**
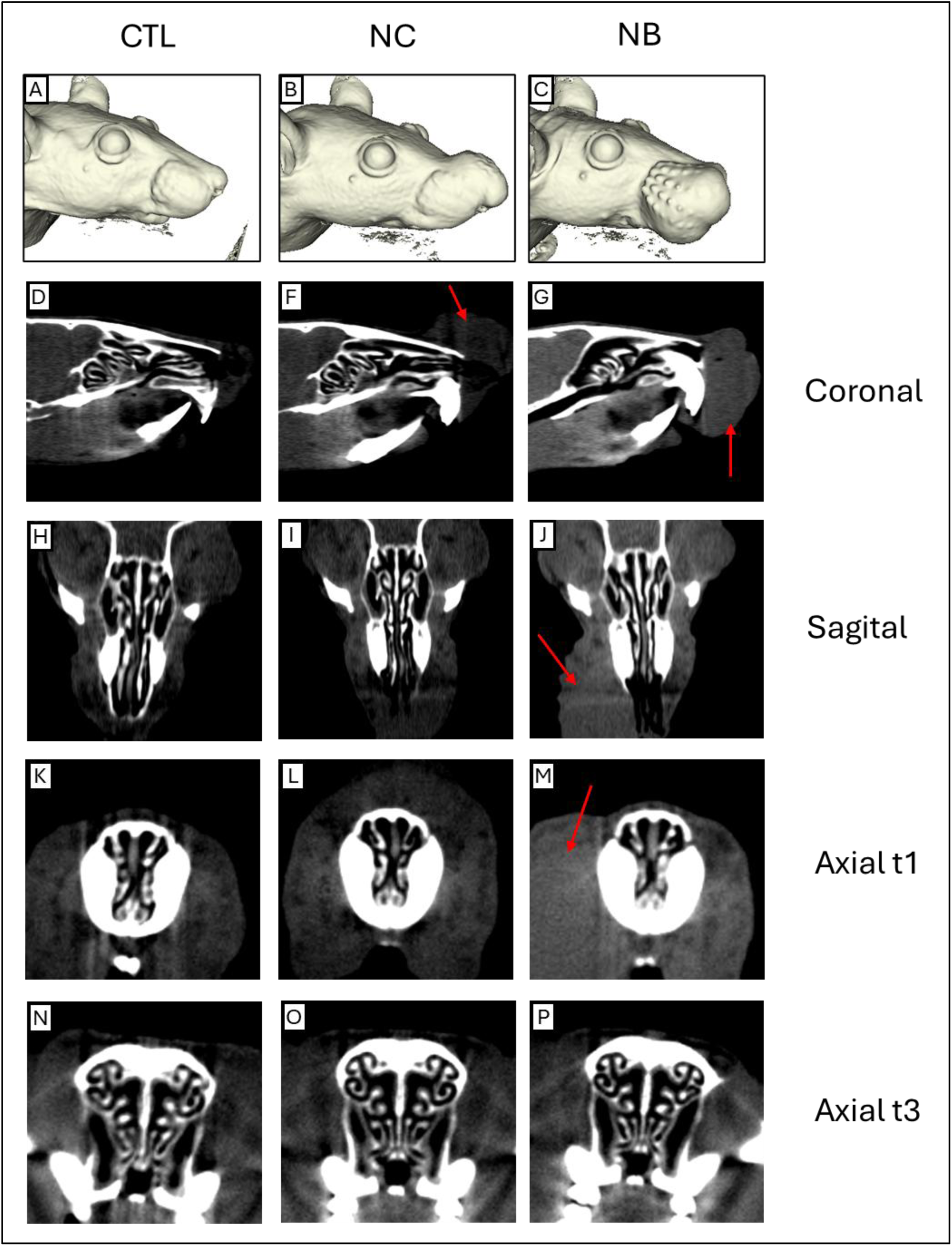
Small animal computed tomography (microCT) The complete three-dimensional reconstructions of the mice (first row). The microCT of the Control (CTL) mock PBS injection in the three sites, Nose Cutaneous (NC) and Nasobasal (NB) groups is also shown, under different section planes: Coronal, Sagittal and Axial at t1 and t3. The red arrows indicate the region of edema. The group NC showed an edema in the nostril region that can progress to the upper part of the snout (second column. While NB infection mode led to a progressive swelling of premaxillary region (third column). No nasal cavity formation was observed, in any of the segments and infection groups.

**Figure S3:**
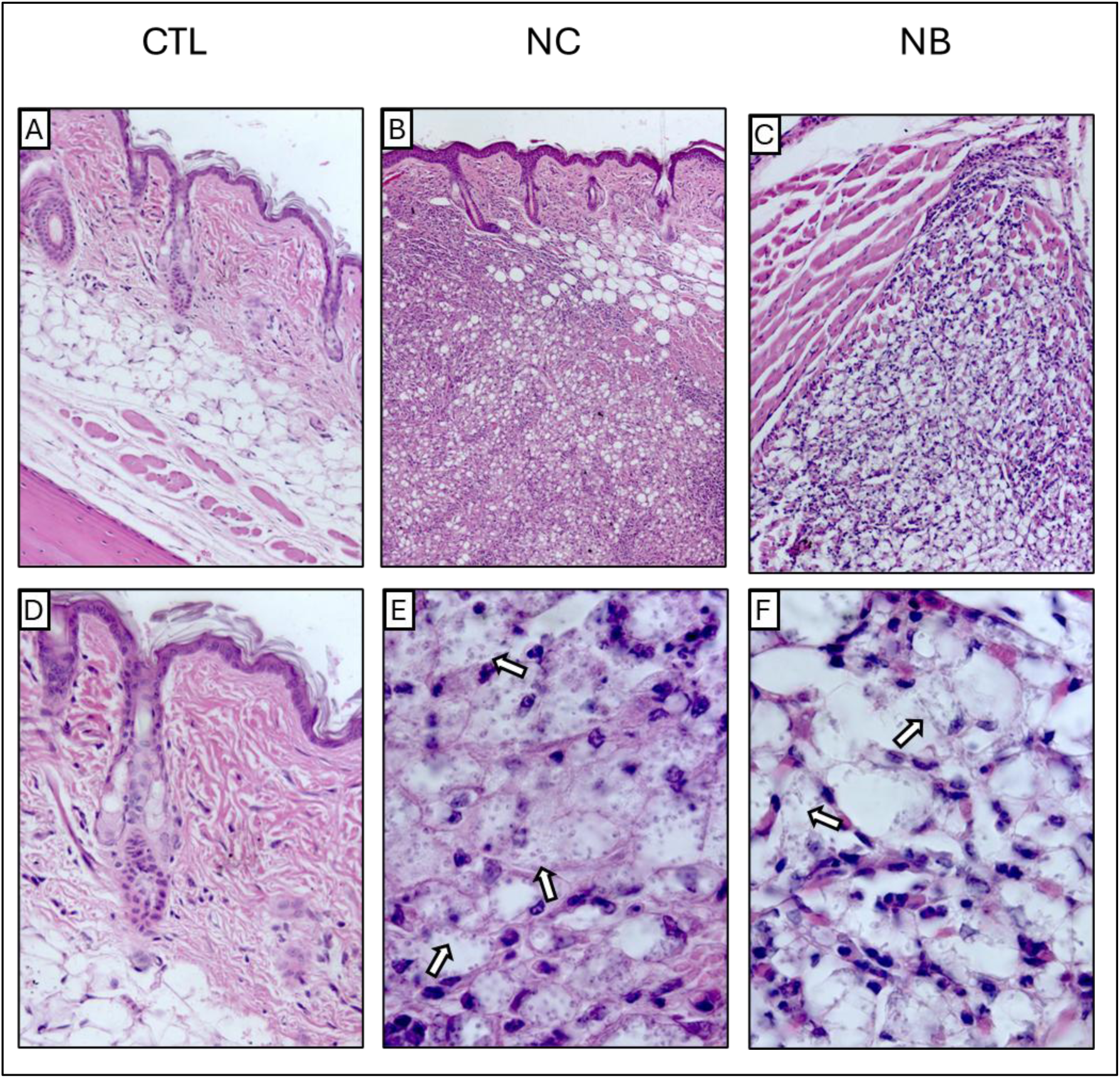
Histopathological Sections of the Epidermal Region of the Nose. Histopathological appearance of the nasal skin tissue from the Control (CTL) mock PBS injection in the three sites, Cutaneous Nose (NC) and Nasobasal (NB) groups, 12weeks after the start of the experiment, stained with hematoxylin and eosin (H&E). The animals were infected with 2×10^6^ parasites. **(A, D)** appearance of the dermis of control animals. **(B, E)** NC group showing inflammatory infiltrate containing infected macrophages, below the epithelium. **(C, F)** NB group showing intense inflammatory infiltrate with parasitized macrophages. Arrows shows examples of macrophages bearing amastigotes. Magnification: (A) 200x; (B) 100x; (C) 100x; (D); 400x; (E) 1000x and (F) 1000x.

**Figure S4:**
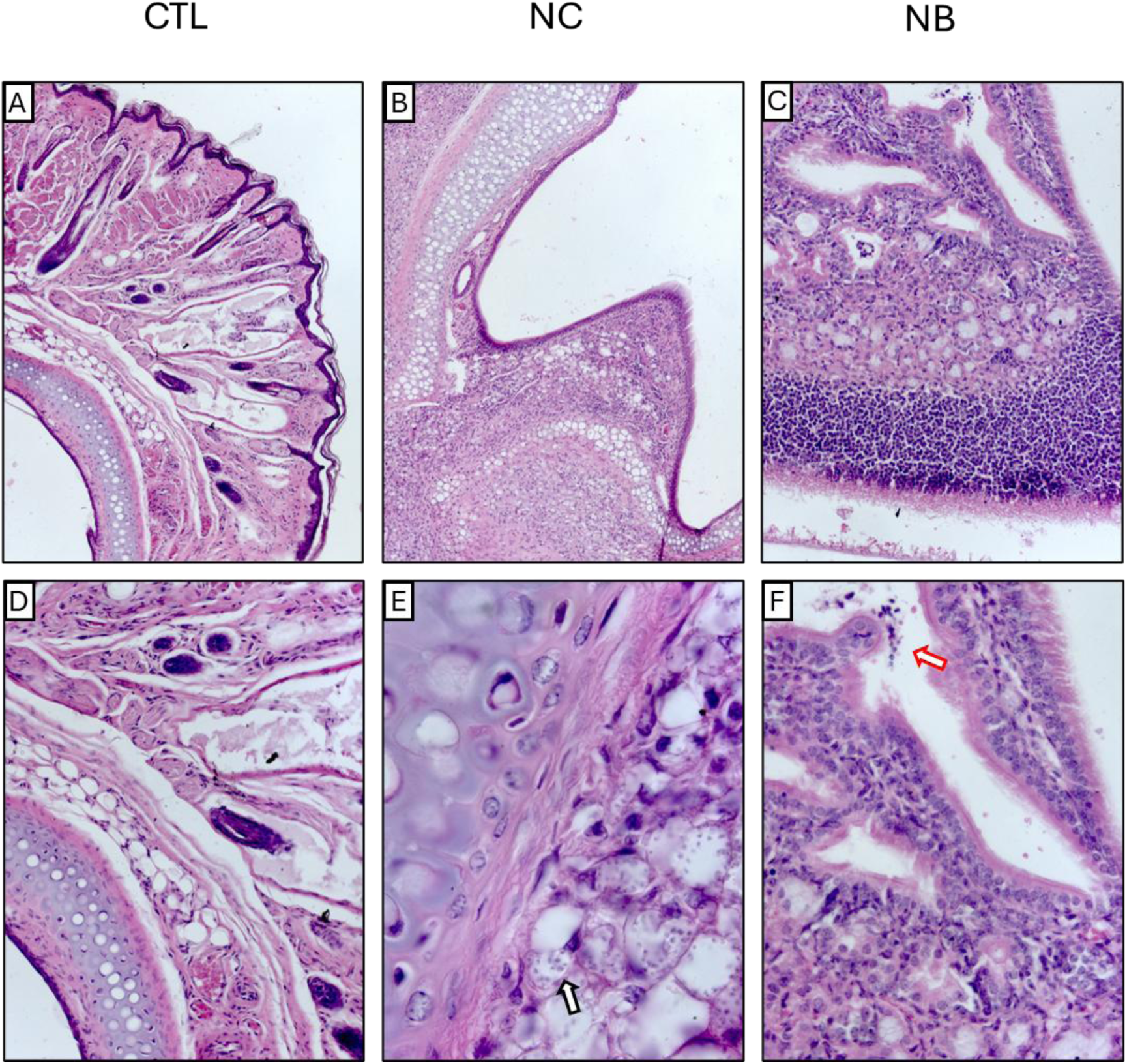
Histopathological Sections of the Initial Internal Region (T1) of the Nasal Mucosa. Histopathological appearance of the nasal tissue from the Control (CTL) mock PBS injection in the three sites, Cutaneous Nose (NC) and Nasobasal (NB) groups, 12weeks after the start of the experiment, stained with hematoxylin and eosin (H&E). The animals were infected with 2×10^6^ parasites. **(A, D)** appearance of the dermis and squamous epithelium of control animals. **(B, E)** NC group showing inflammatory infiltrate containing infected macrophages, close to the cartilaginous regions. **(C, F)** NB group, demonstrating stress and structural disorganization of the tissue accompanied by cell detachment. White arrows show examples of macrophages bearing amastigotes, red arrows show example of cell detachment. Magnification: (A) 100x; (B) 100x; (C) 100x; (D); 200x; (E) 1000x and (F) 200x.

**Figure S5.**
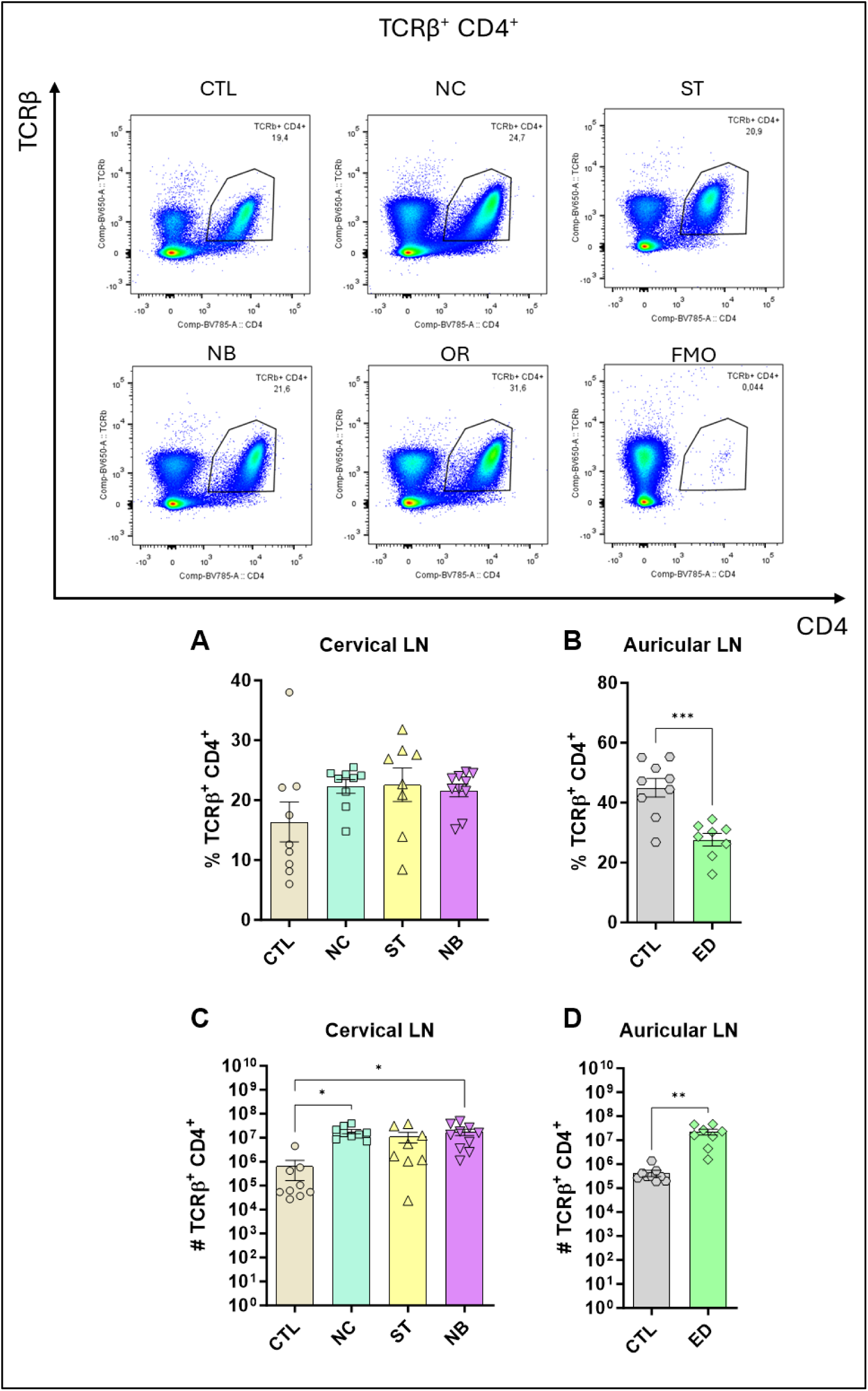
Profile of TCRβ^+^ CD4^+^ Cells. The figure shows the representatives and graphs with the percentage of cells in the nasal and ear sites (A) and (B) respectively. In (C) and (D) data on the total number of cells are represented. Data accumulative of two independent experiments. Statistics: plot with Standard Error of The Mean (SEM), t-test was used for all groups and samples * p<0,05, **<0,005., 3-6 animals per group in each.

**Figure S6.**
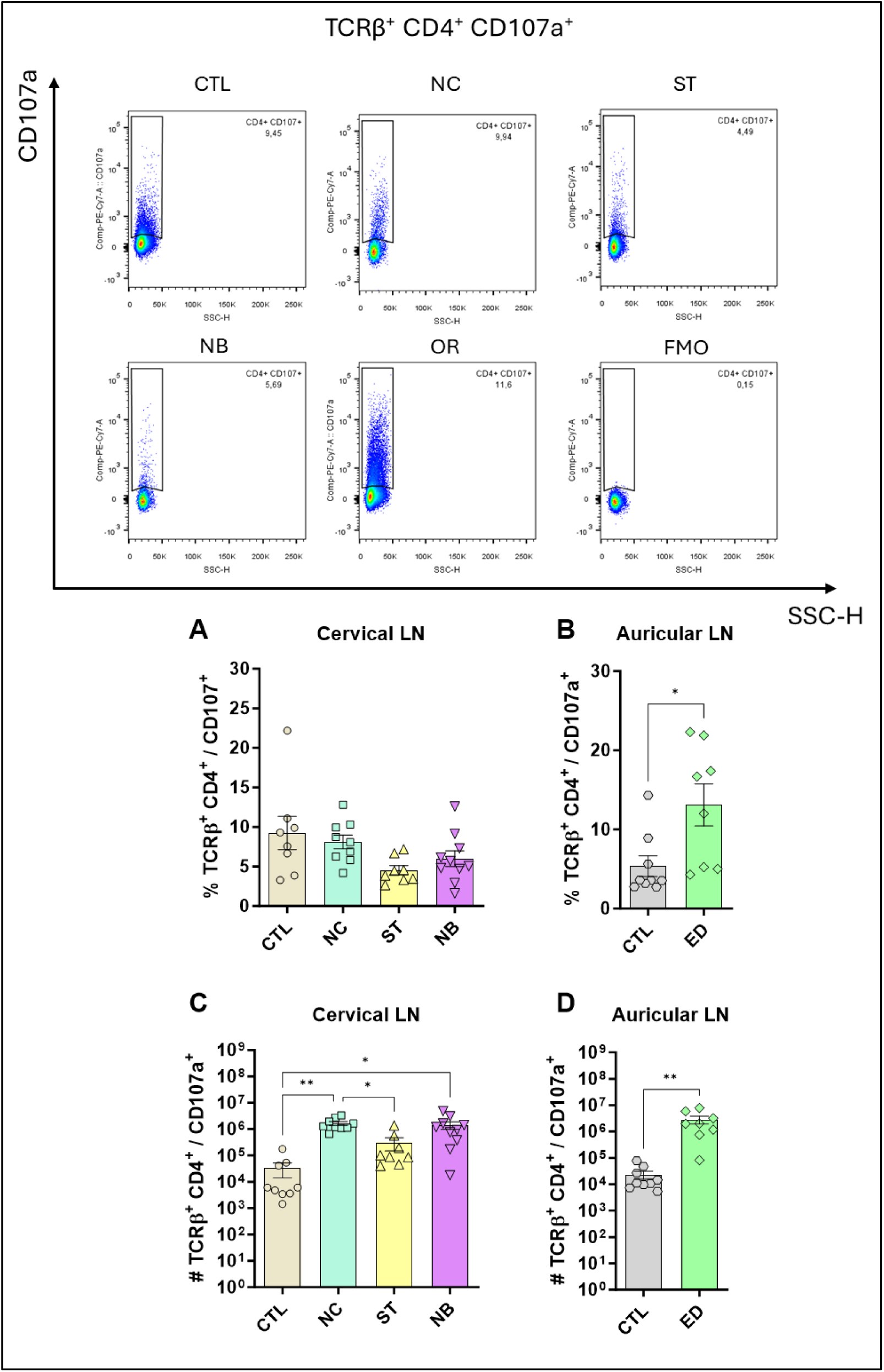
Profile of TCRβ^+^ CD4^+^ CD107a^+^ Cells. The figure shows the representatives and graphs with the percentage of cells in the nasal and ear sites (A) and (B) respectively. In (C) and (D) data on the total number of cells are represented. Data accumulative of two independent experiments. Statistics: plot with Standard Error of The Mean (SEM), t-test was used for all groups and samples * p<0,05, **<0,005., 3-6 animals per group in each.

**Figure S7.**
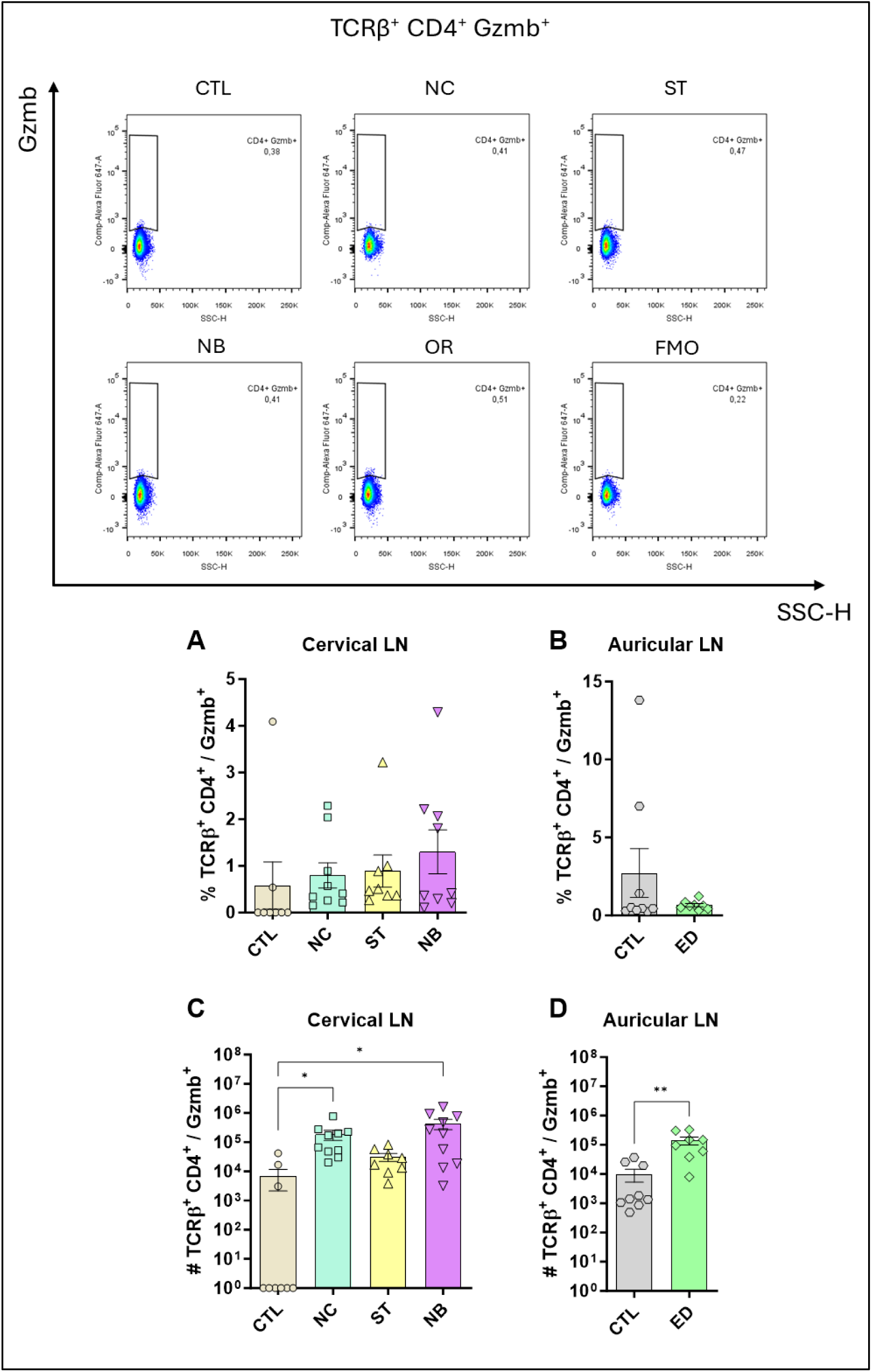
Profile of TCRβ^+^ CD4^+^ Gzmb^+^ Cells. The figure shows the representatives and graphs with the percentage of cells in the nasal and ear sites (A) and (B) respectively. In (C) and (D) data on the total number of cells are represented. Data accumulative of two independent experiments. Statistics: plot with Standard Error of The Mean (SEM), t-test was used for all groups and samples * p<0,05, **<0,005., 3-6 animals per group in each.

**Figure S8.**
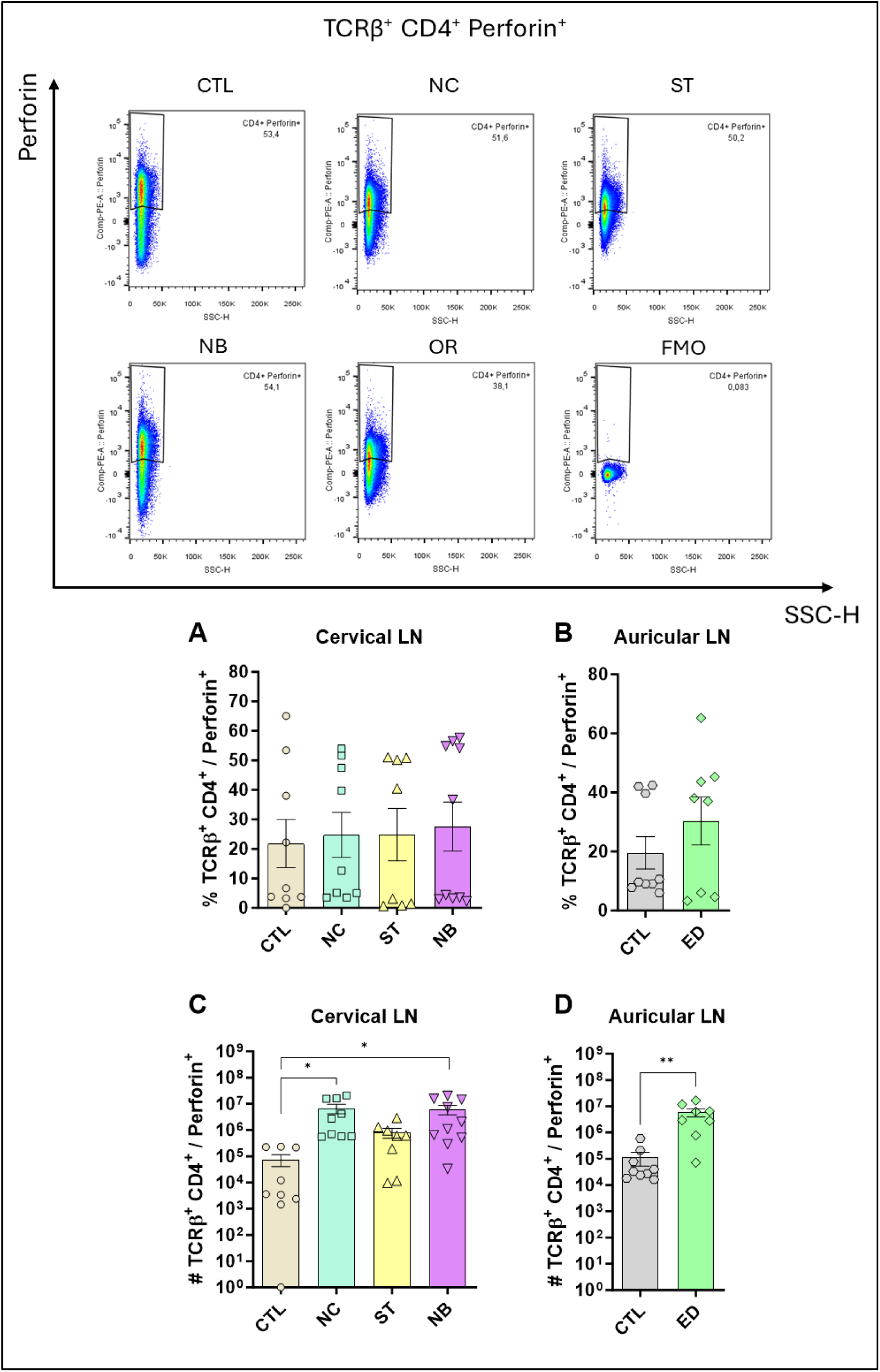
Profile of TCRβ^+^ CD4^+^ Perforin^+^ Cells. The figure shows the representatives and graphs with the percentage of cells in the nasal and ear sites (A) and (B) respectively. In (C) and (D) data on the total number of cells are represented. Data accumulative of two independent experiments. Statistics: plot with Standard Error of The Mean (SEM), t-test was used for all groups and samples * p<0,05, **<0,005., 3-6 animals per group in each.

**Figure S9.**
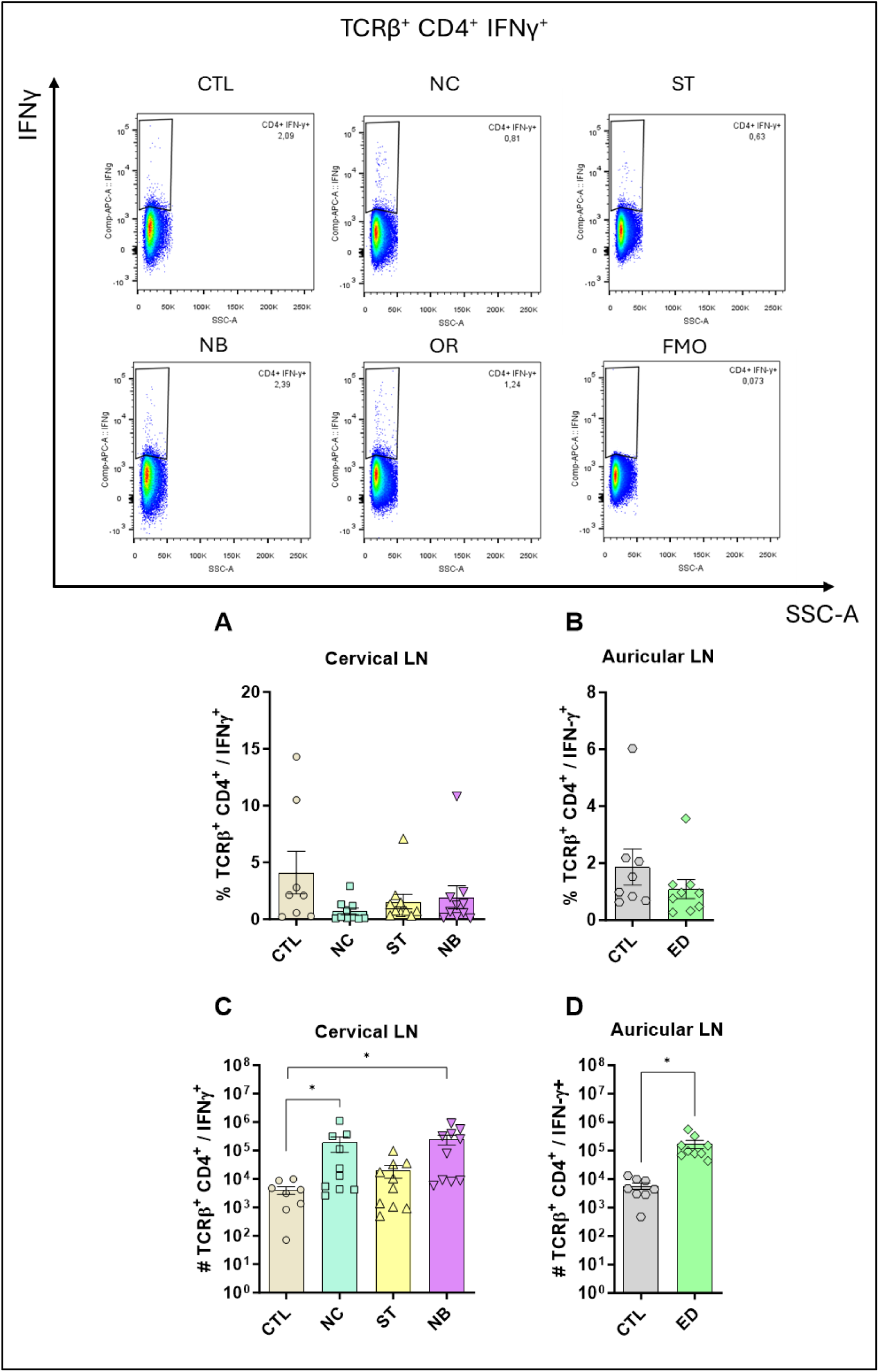
Profile of TCRβ^+^ CD4^+^ IFN-γ^+^ Cells. The figure shows the representatives and graphs with the percentage of cells in the nasal and ear sites (A) and (B) respectively. In (C) and (D) data on the total number of cells are represented. Data accumulative of two independent experiments. Statistics: plot with Standard Error of The Mean (SEM), t-test was used for all groups and samples * p<0,05, **<0,005., 3-6 animals per group in each.

**Figure S10.**
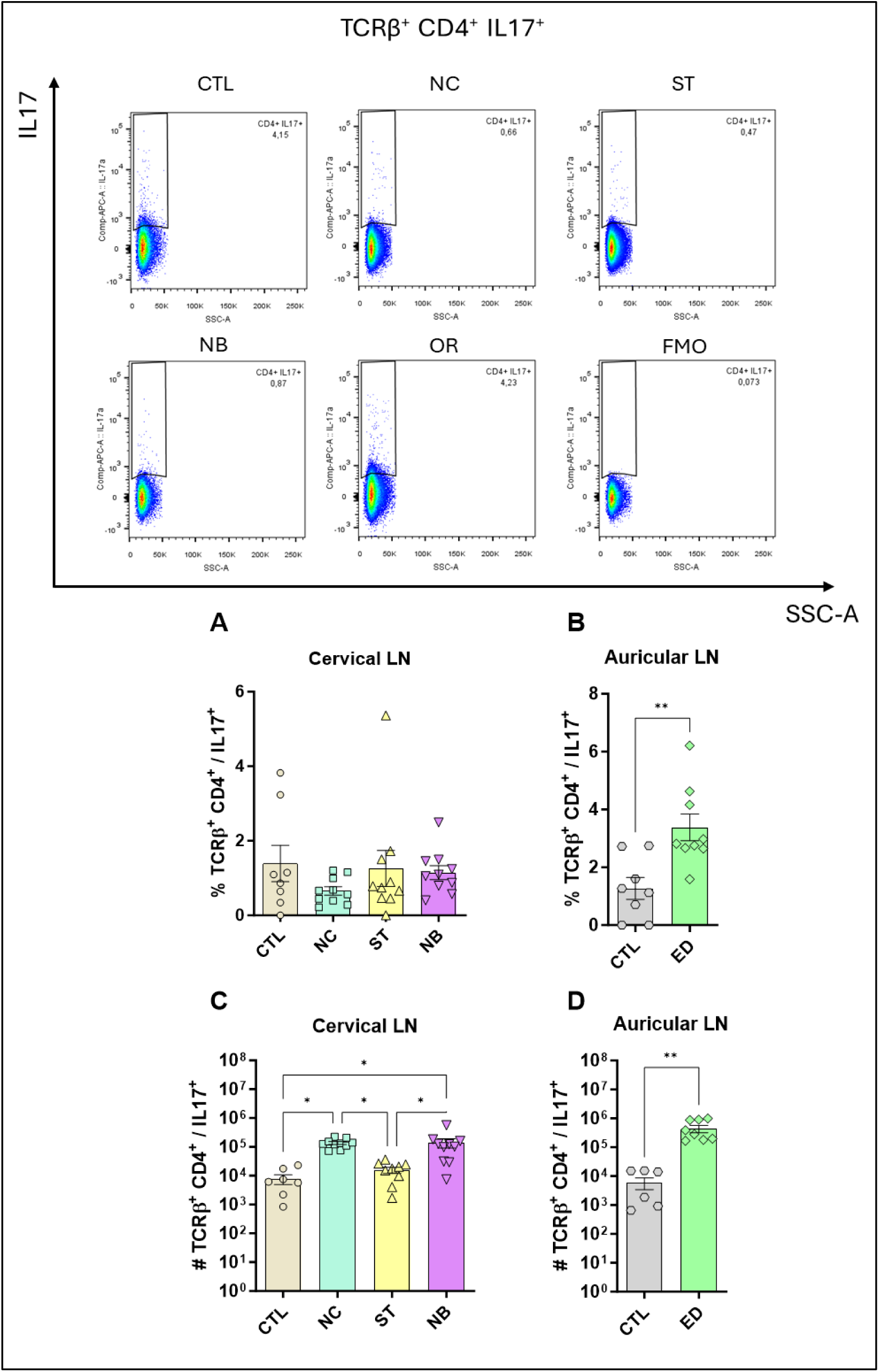
Profile of TCRβ^+^ CD4^+^ IL-17^+^ Cells. The figure shows the representatives and graphs with the percentage of cells in the nasal and ear sites (A) and (B) respectively. In (C) and (D) data on the total number of cells are represented. Data accumulative of two independent experiments. Statistics: plot with Standard Error of The Mean (SEM), t-test was used for all groups and samples * p<0,05, **<0,005., 3-6 animals per group in each.

**Figure S11.**
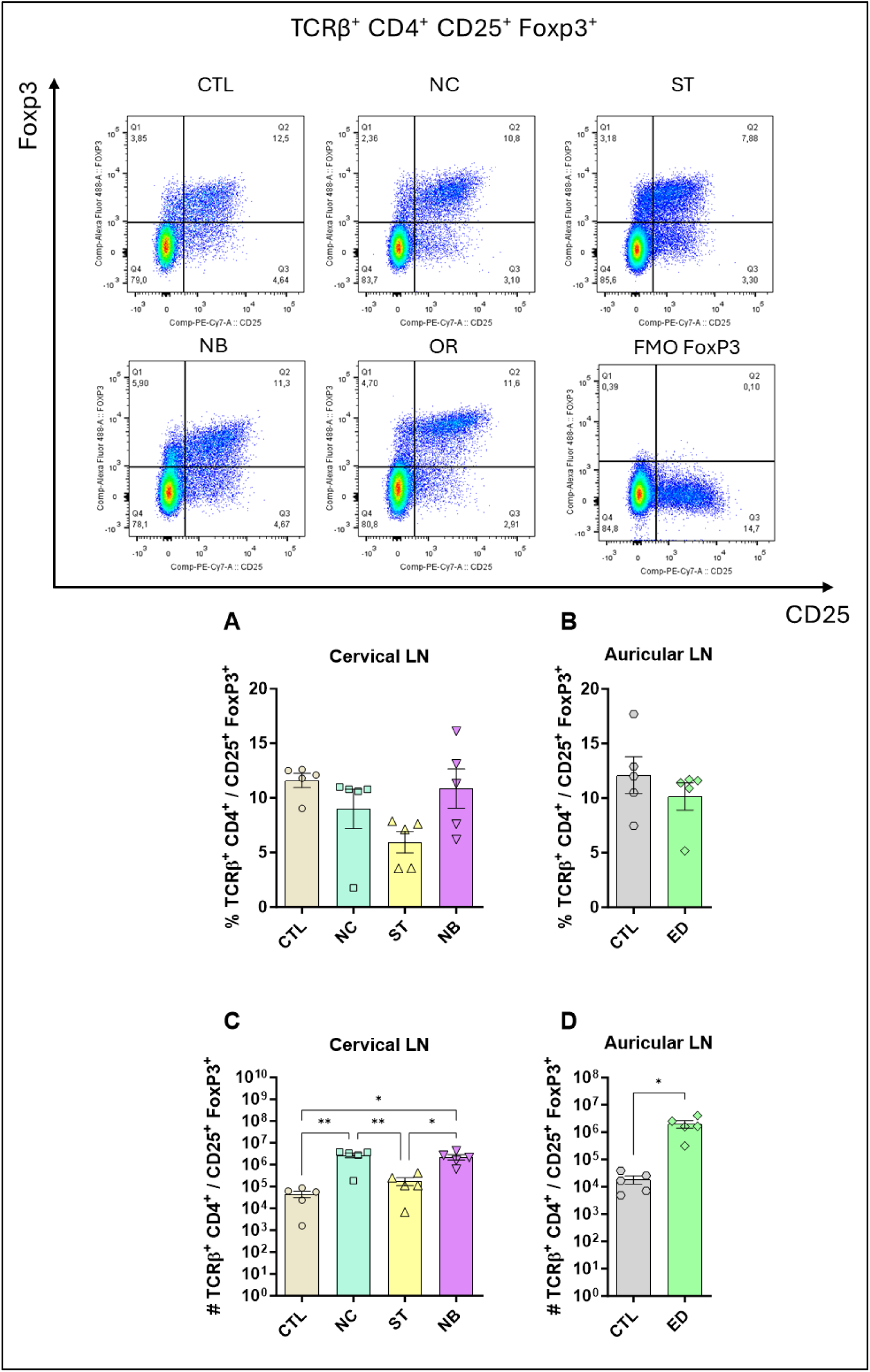
TCRβ^+^ CD4^+^ CD25^-^ FoxP3^+^ profile. The figure shows the representatives and graphs with the percentage of cells in the nasal and ear sites (A) and (B) respectively. In (C) and (D) data on the total number of cells are represented. Data accumulative of two independent experiments. Statistics: plot with Standard Error of The Mean (SEM), t-test was used for all groups and samples * p<0,05, **<0,005., 3-6 animals per group in each.

**Figure S12.**
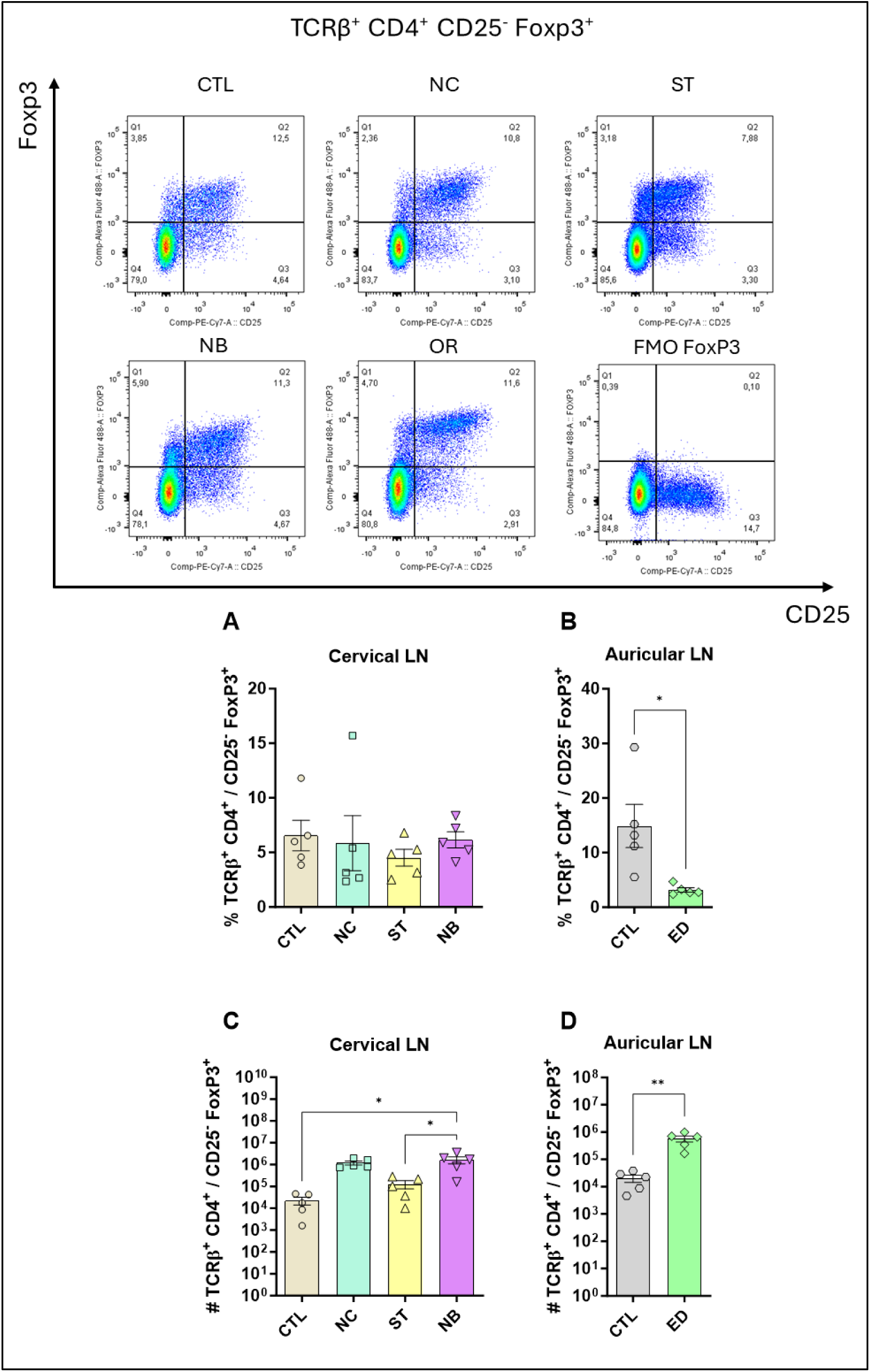
TCRβ^+^ CD4^+^ CD25^-^ FoxP3^+^ profile. The figure shows the representatives and graphs with the percentage of cells in the nasal and ear sites (A) and (B) respectively. In (C) and (D) data on the total number of cells are represented. Data of one independent experiment. Statistics: plot with Standard Error of The Mean (SEM), t-test was used for all groups and samples p<0,05, **<0,005., 3-6 animals per group in each.

**Figure S13.**
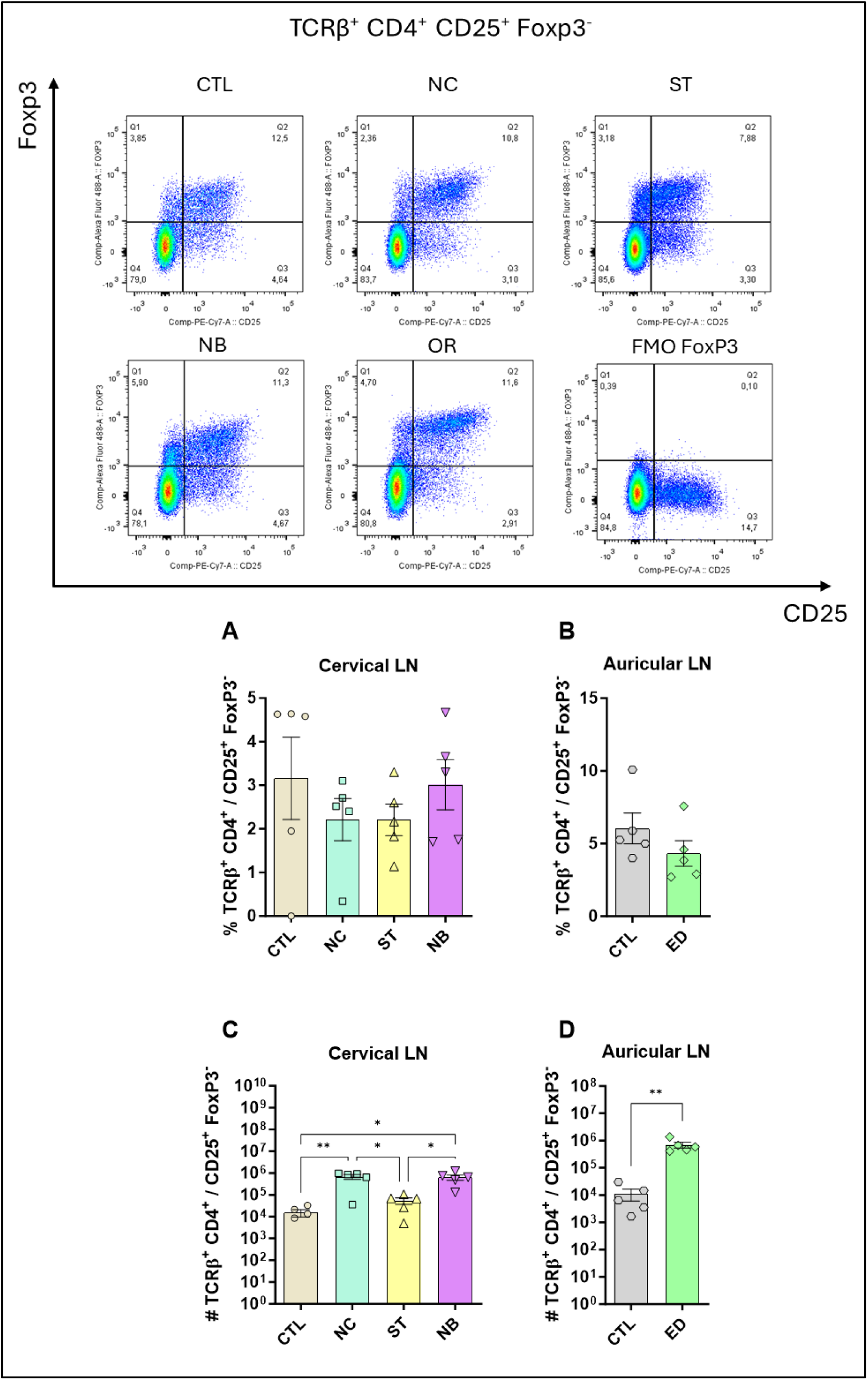
TCRβ^+^ CD4^+^ CD25^+^ FoxP3^-^ profile. The figure shows the representatives and graphs with the percentage of cells in the nasal and ear sites (A) and (B) respectively. In (C) and (D) data on the total number of cells are represented. Data of one independent experiment. Statistics: plot with Standard Error of The Mean (SEM), t-test was used for all groups and samples p<0,05, **<0,005., 3-6 animals per group in each.

**Figure S14.**
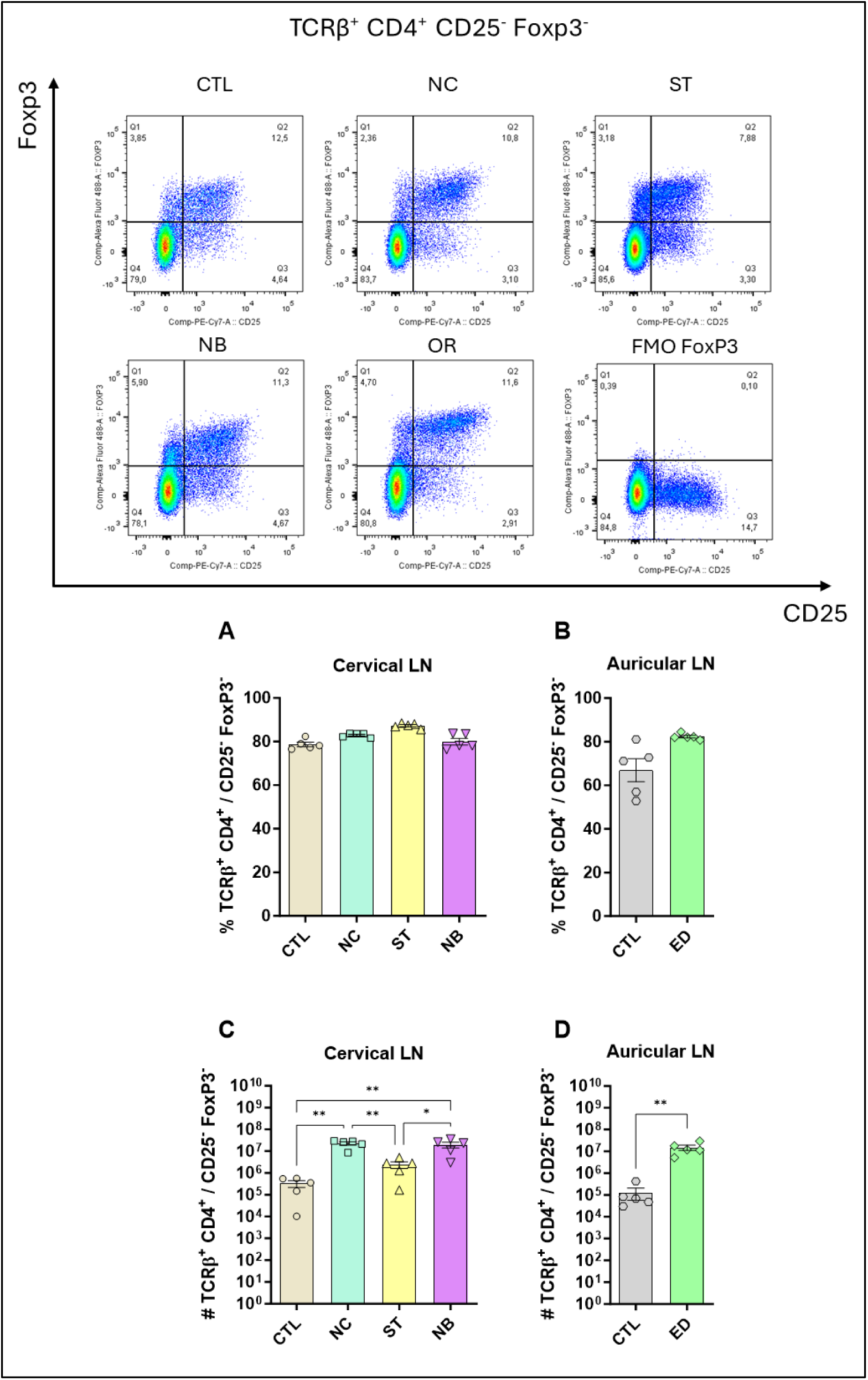
TCRβ^+^ CD4^+^ CD25^-^ FoxP3^-^ profile. The figure shows the representatives and graphs with the percentage of cells in the nasal and ear sites (A) and (B) respectively. In (C) and (D) data on the total number of cells are represented. Data of one independent experiment. Statistics: plot with Standard Error of The Mean (SEM), t-test was used for all groups and samples p<0,05, **<0,005., 3-6 animals per group in each.

**Figure S15.**
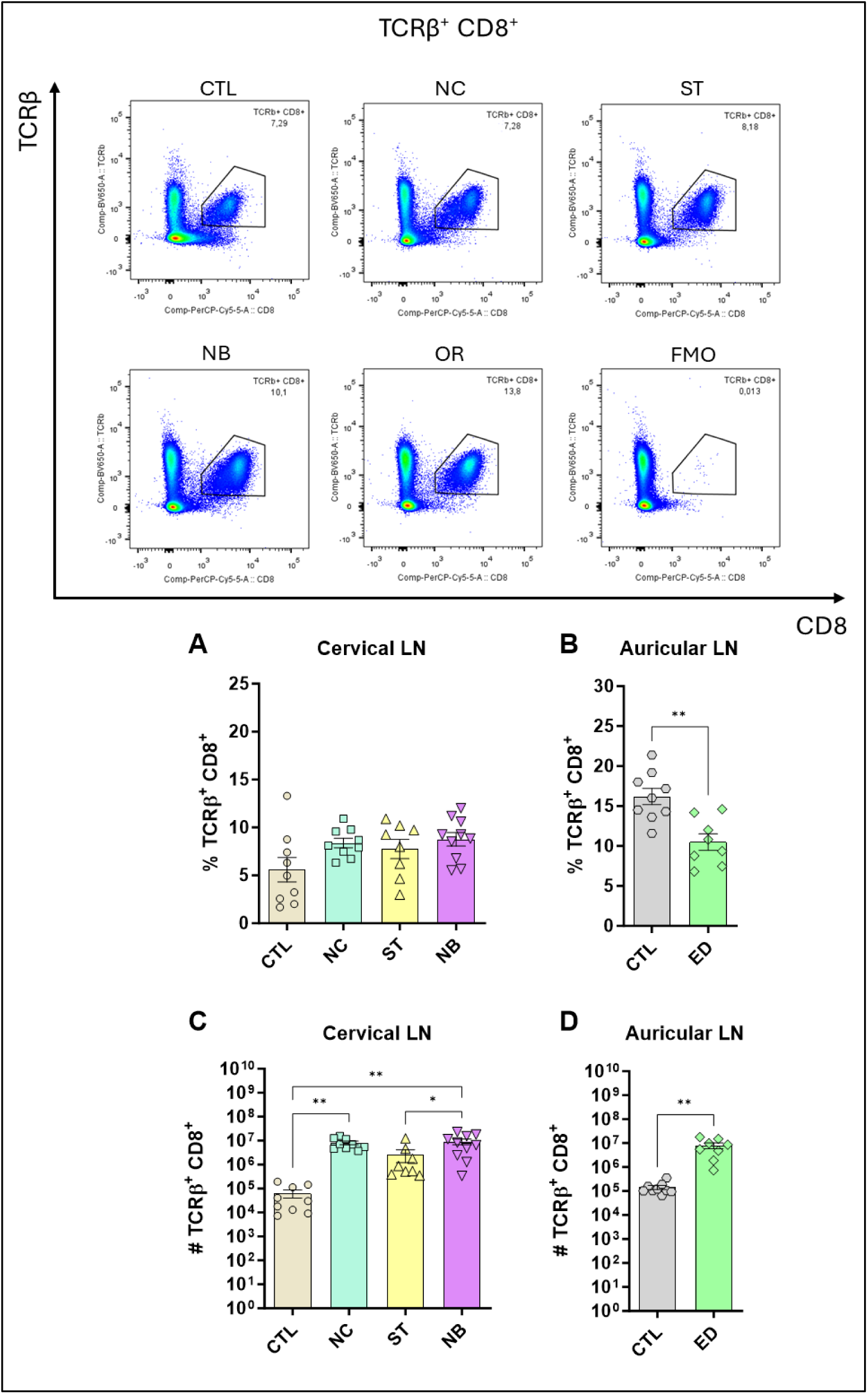
Profile of TCRβ^+^ CD8^+^ Cells. The figure shows the representatives and graphs with the percentage of cells in the nasal and ear sites (A) and (B) respectively. In (C) and (D) data on the total number of cells are represented. Data accumulative of two independent experiments. Statistics: plot with Standard Error of The Mean (SEM), t-test was used for all groups and samples p<0,05, **<0,005., 3-6 animals per group in each.

**Figure S16.**
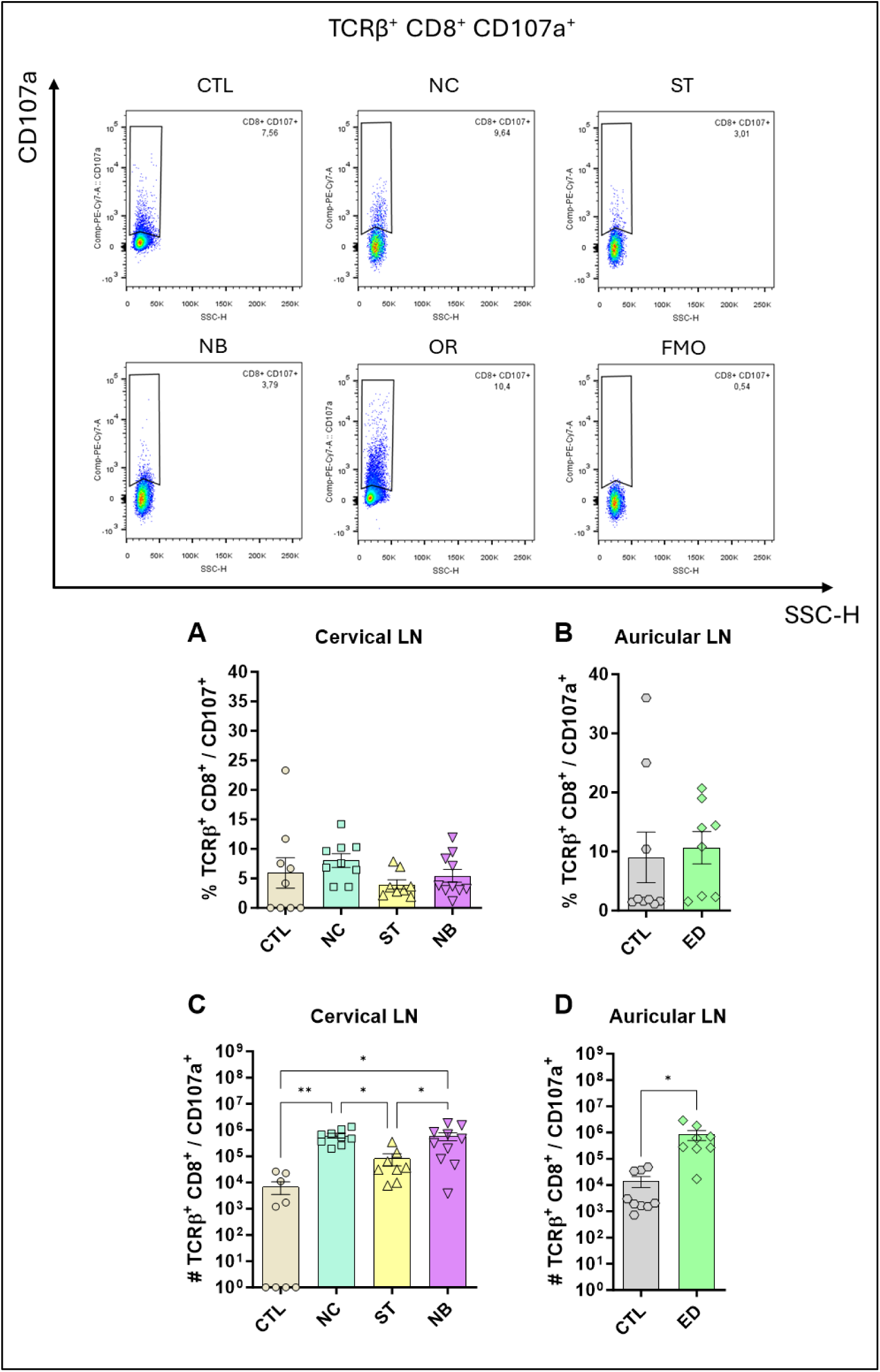
Profile of TCRβ^+^ CD8^+^ CD107a^+^ Cells. The figure shows the representatives and graphs with the percentage of cells in the nasal and ear sites (A) and (B) respectively. In (C) and (D) data on the total number of cells are represented. Data accumulative of two independent experiments. Statistics: plot with Standard Error of The Mean (SEM), t-test was used for all groups and samples * p<0,05, **<0,005., 3-6 animals per group in each.

**Figure S17.**
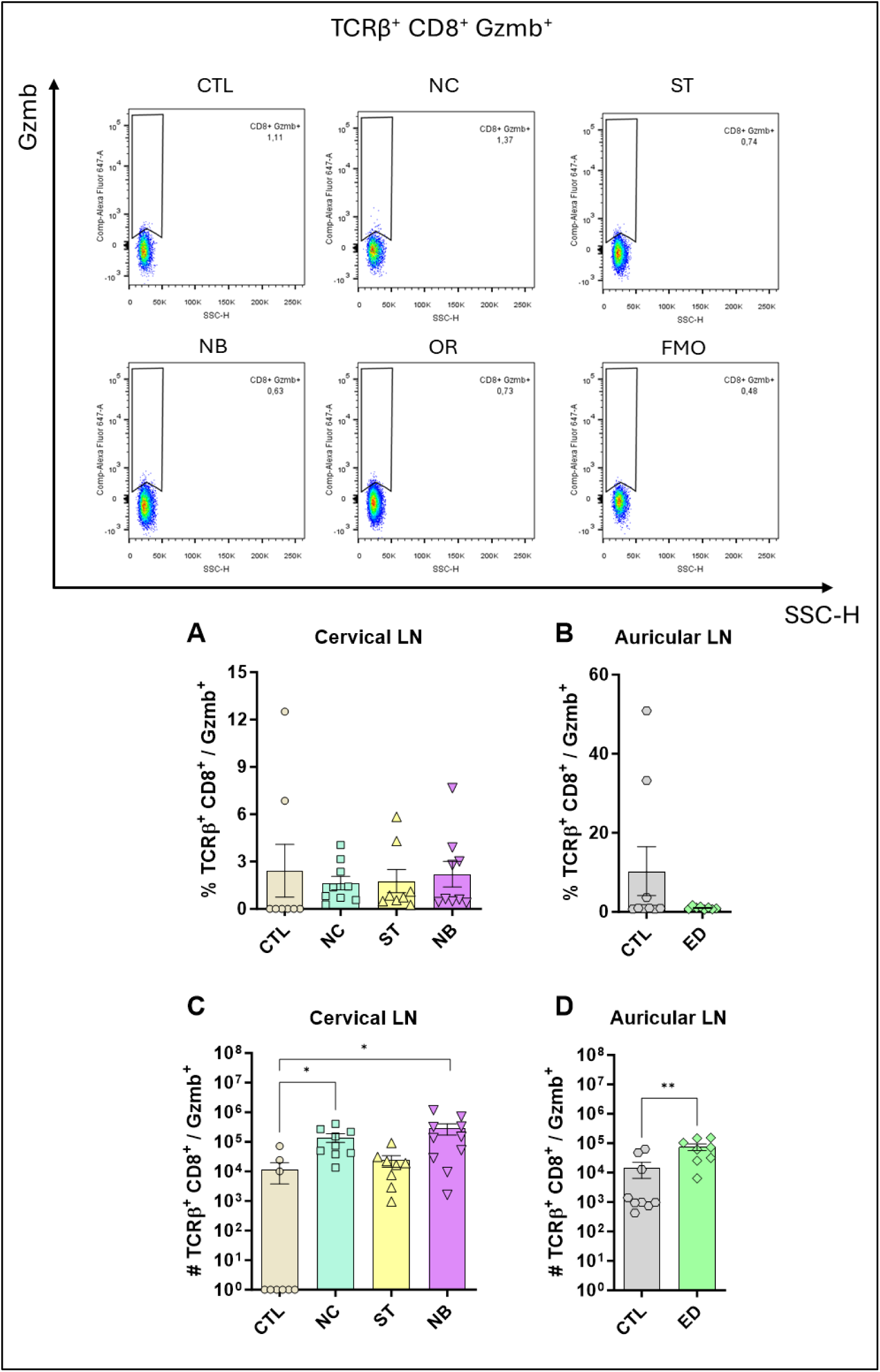
Profile of TCRβ^+^ CD8^+^ Gzmb^+^ Cells. The figure shows the representatives and graphs with the percentage of cells in the nasal and ear sites (A) and (B) respectively. In (C) and (D) data on the total number of cells are represented. Data accumulative of two independent experiments. Statistics: plot with Standard Error of The Mean (SEM), t-test was used for all groups and samples * p<0,05, **<0,005., 3-6 animals per group in each.

**Figure S18.**
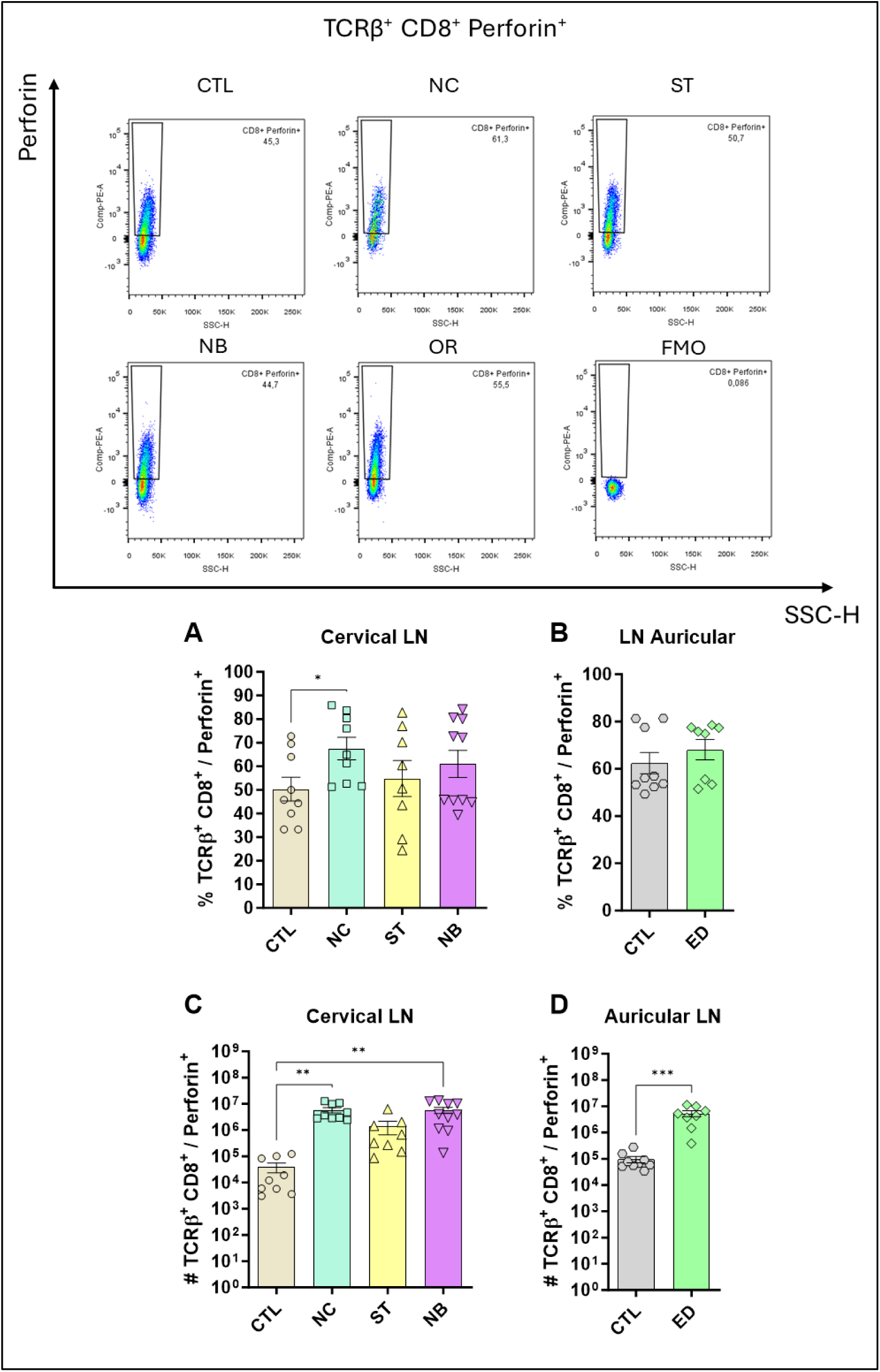
Profile of TCRβ^+^ CD8^+^ Perforin^+^ Cells. The figure shows the representatives and graphs with the percentage of cells in the nasal and ear sites (A) and (B) respectively. In (C) and (D) data on the total number of cells are represented. Data accumulative of two independent experiments. Statistics: plot with Standard Error of The Mean (SEM), t-test was used for all groups and samples * p<0,05, **<0,005., 3-6 animals per group in each.

**Figure S19.**
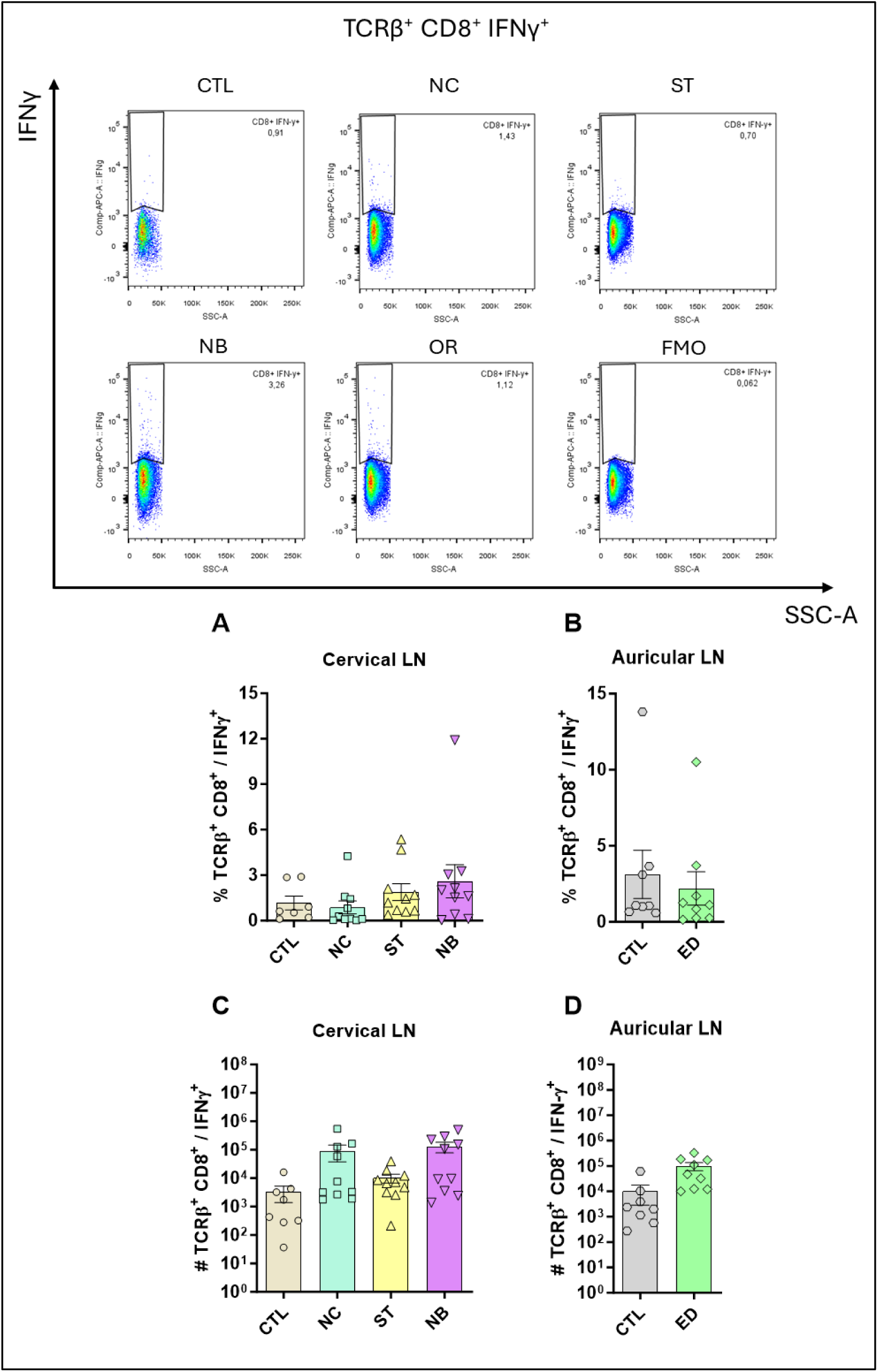
Profile of TCRβ^+^ CD8^+^ IFN-γ^+^ Cells. The figure shows the representatives and graphs with the percentage of cells in the nasal and ear sites (A) and (B) respectively. In (C) and (D) data on the total number of cells are represented. Data accumulative of two independent experiments. Statistics: plot with Standard Error of The Mean (SEM), t-test was used for all groups and samples * p<0,05, **<0,005., 3-6 animals per group in each.

**Figure S20.**
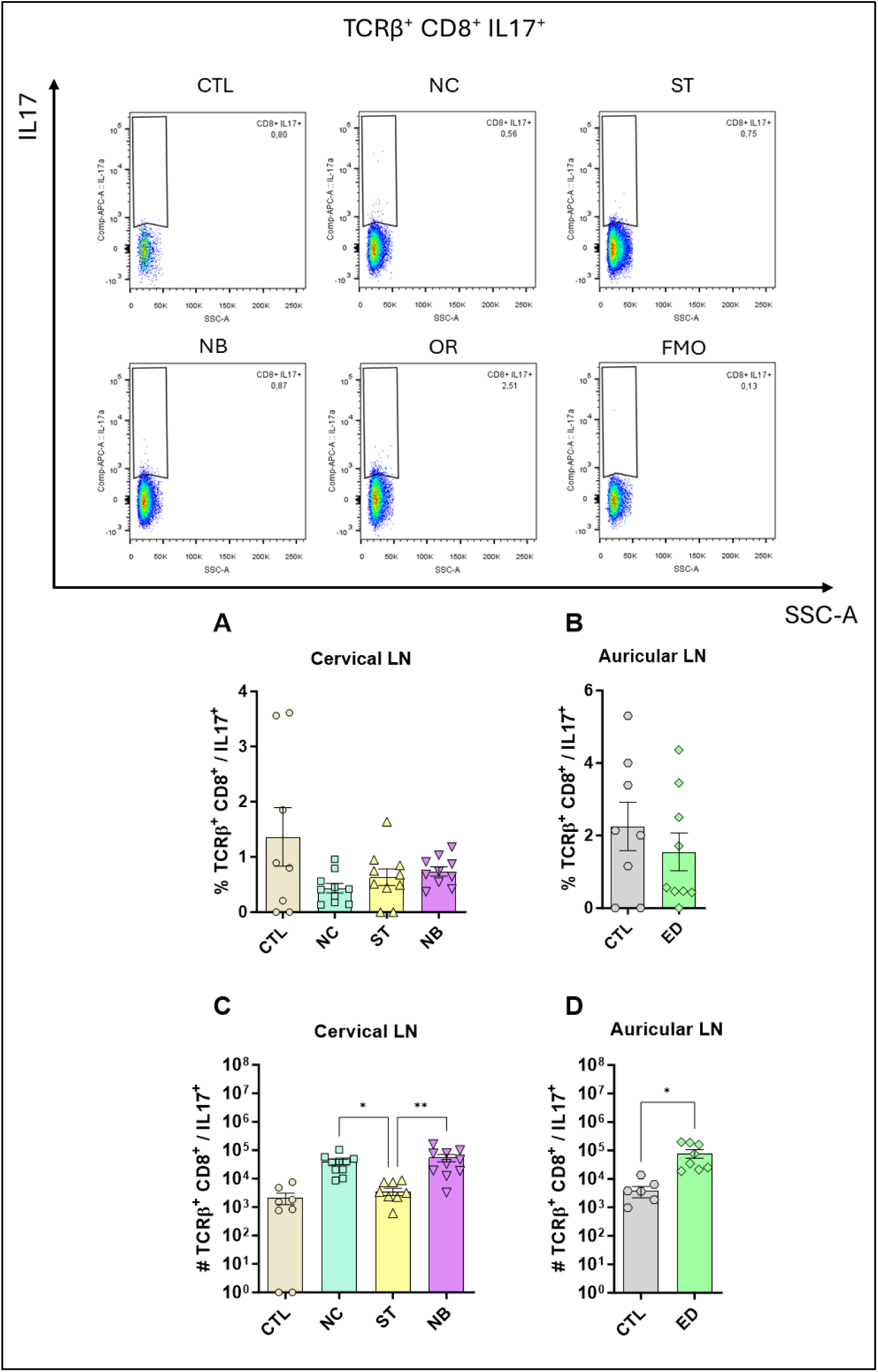
Profile of TCRβ^+^ CD8^+^ IL-17^+^ Cells. The figure shows the representatives and graphs with the percentage of cells in the nasal and ear sites (A) and (B) respectively. In (C) and (D) data on the total number of cells are represented. Data accumulative of two independent experiments. Statistics: plot with Standard Error of The Mean (SEM), t-test was used for all groups and samples * p<0,05, **<0,005., 3-6 animals per group in each.

**Figure S21.**
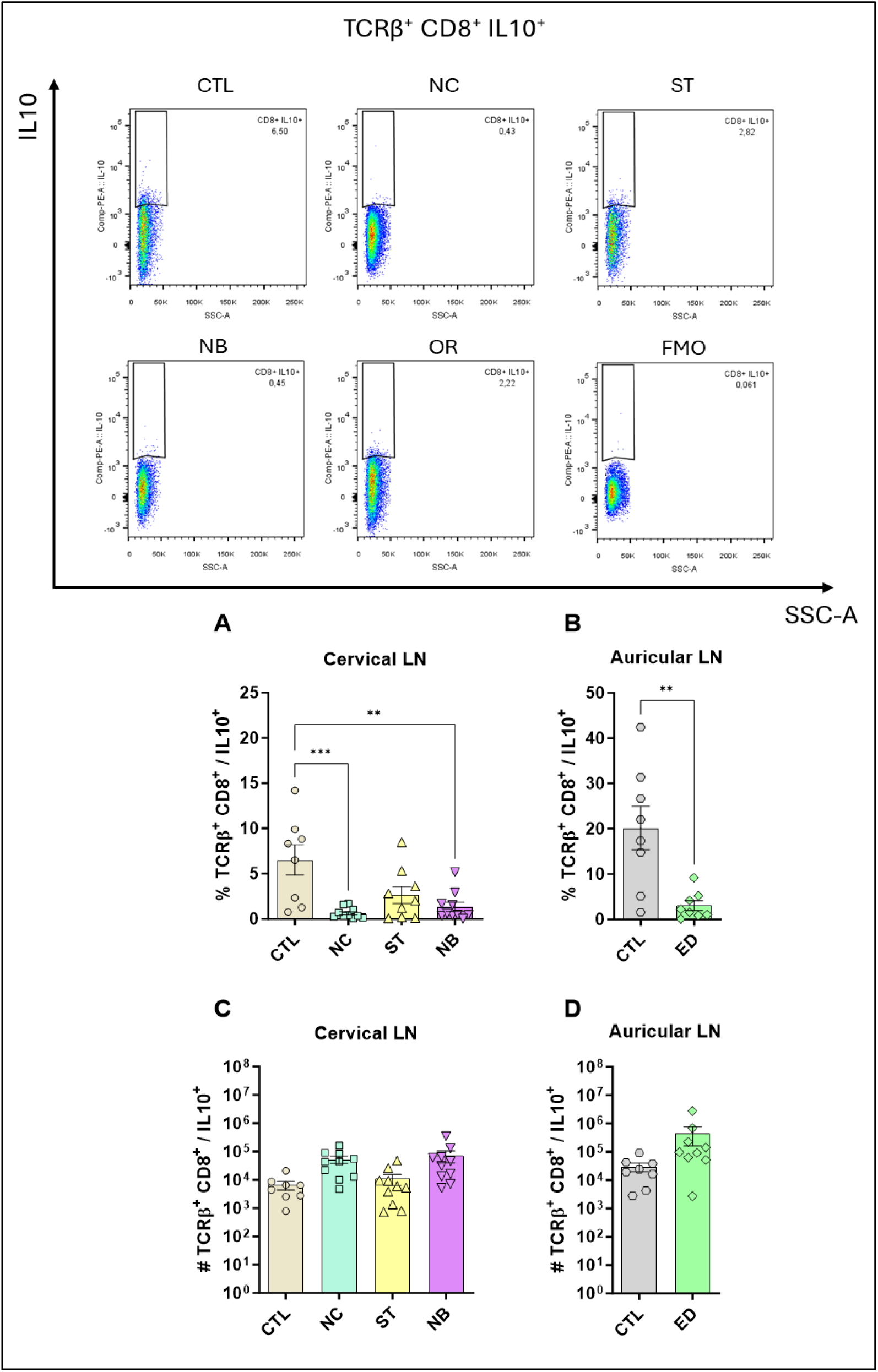
Profile of TCRβ^+^ CD8^+^ IL-10^+^ Cells. The figure shows the representatives and graphs with the percentage of cells in the nasal and ear sites (A) and (B) respectively. In (C) and (D) data on the total number of cells are represented. Data accumulative of two independent experiments. Statistics: plot with Standard Error of The Mean (SEM), t-test was used for all groups and samples * p<0,05, **<0,005., 3-6 animals per group in each.

**Figure S22.**
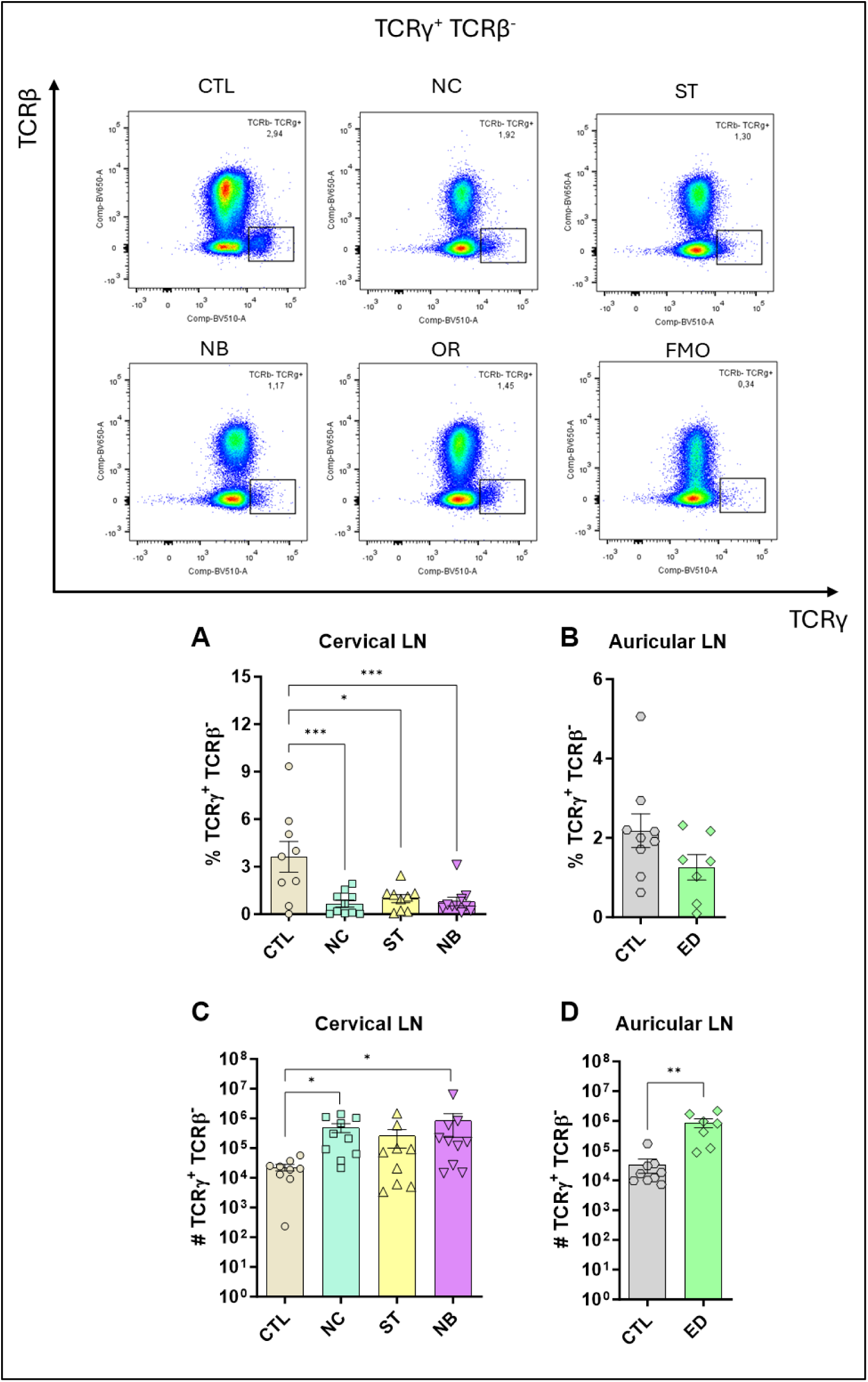
Profile of TCRγ^+^ TCRβ^-^ Cells. The figure shows the representatives and graphs with the percentage of cells in the nasal and ear sites (A) and (B) respectively. In (C) and (D) data on the total number of cells are represented. Data accumulative of two independent experiments. Statistics: plot with Standard Error of The Mean (SEM), t-test was used for all groups and samples p<0,05, **<0,005., 3-6 animals per group in each.

**Figure S23.**
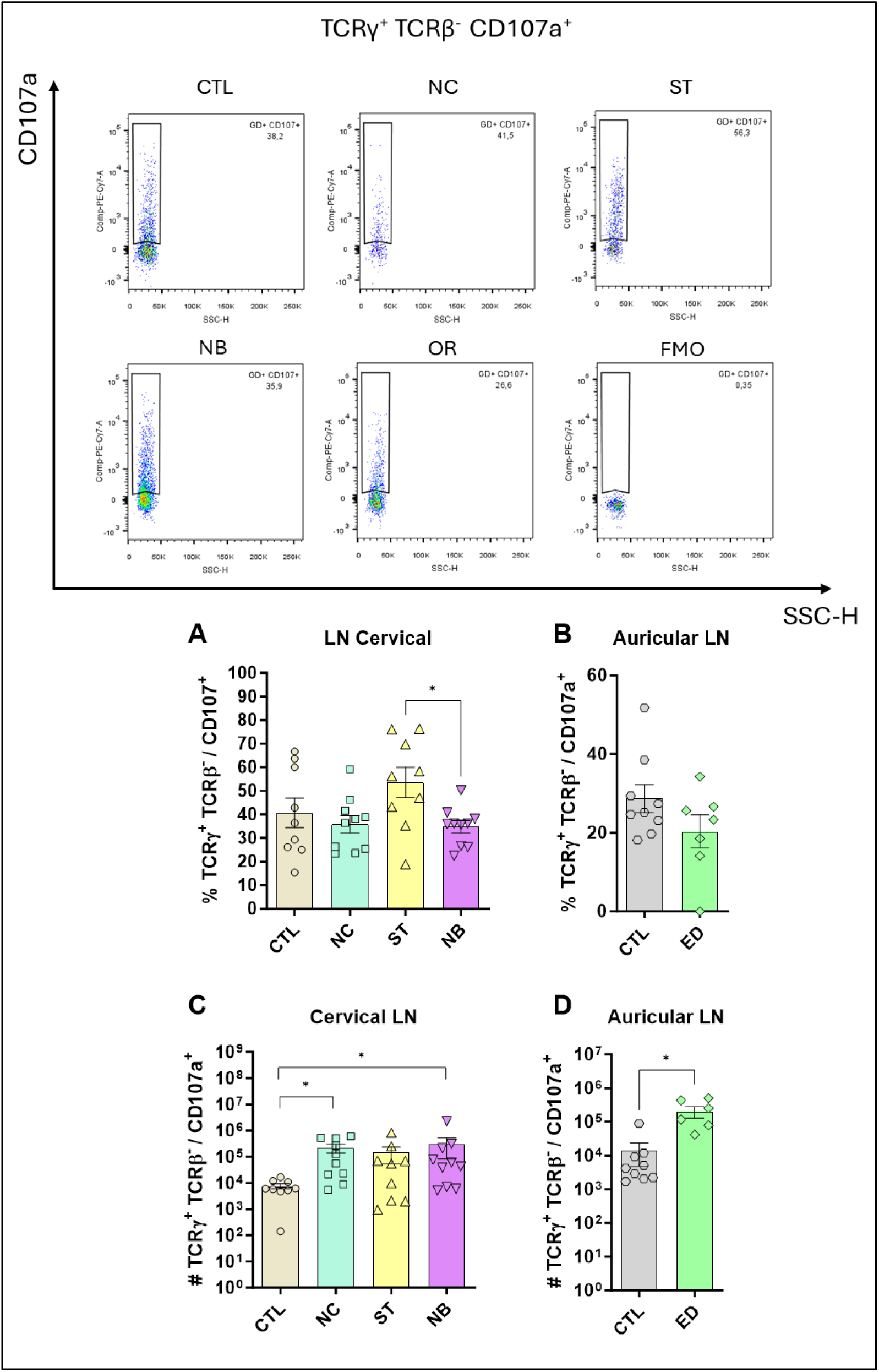
TCRγ^+^ TCRβ^-^ CD107a^+^ profile. The figure shows the representatives and graphs with the percentage of cells in the nasal and ear sites (A) and (B) respectively. In (C) and (D) data on the total number of cells are represented. Data accumulative of two independent experiments. Statistics: plot with Standard Error of The Mean (SEM), t-test was used for all groups and samples p<0,05, **<0,005., 3-6 animals per group in each.

**Figure S24.**
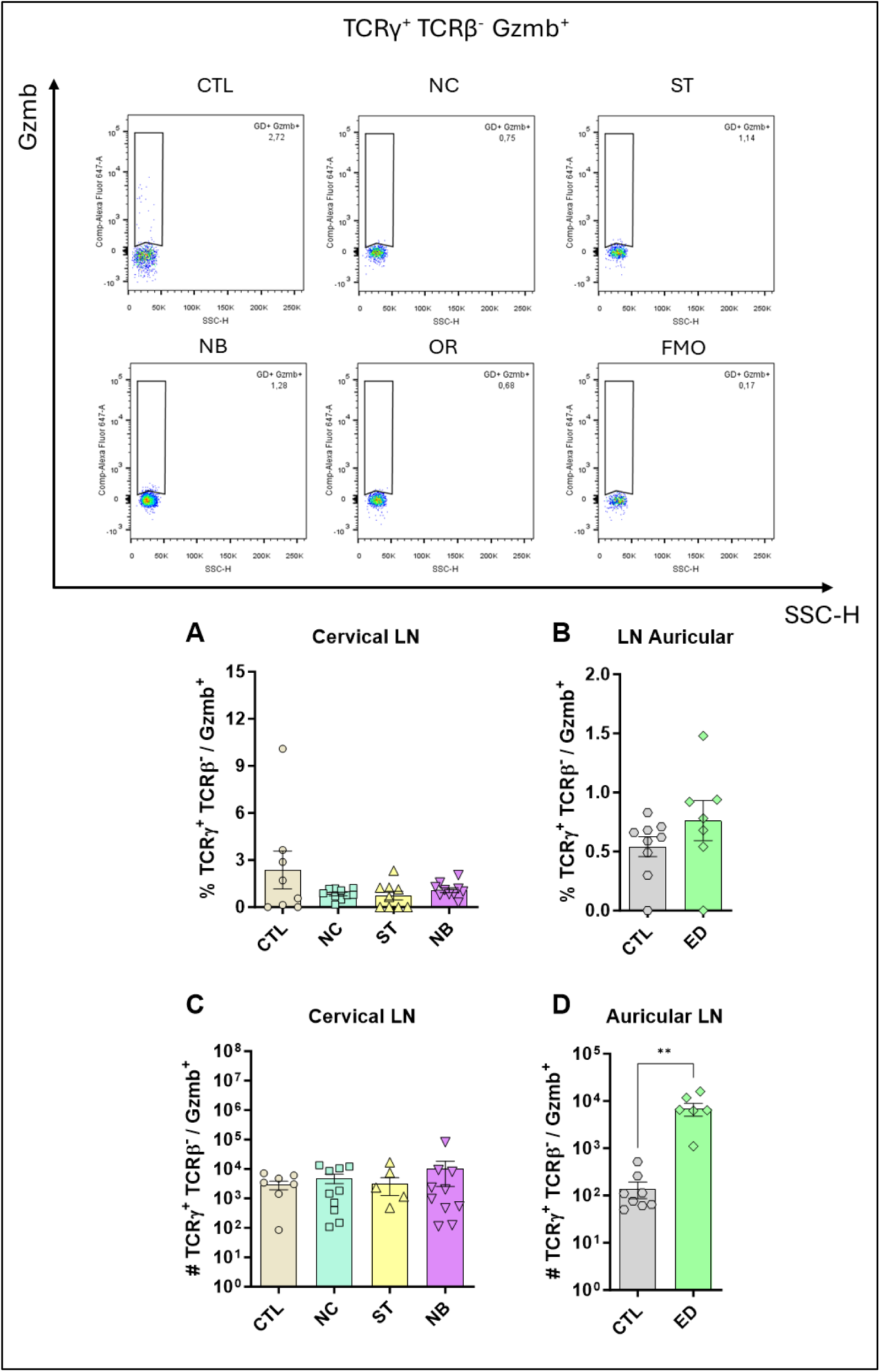
TCRγ^+^ TCRβ^-^ Gzmb^+^ profile. The figure shows the representatives and graphs with the percentage of cells in the nasal and ear sites (A) and (B) respectively. In (C) and (D) data on the total number of cells are represented. Data accumulative of two independent experiments. Statistics: plot with Standard Error of The Mean (SEM), t-test was used for all groups and samples p<0,05, **<0,005., 3-6 animals per group in each.

**Figure S25.**
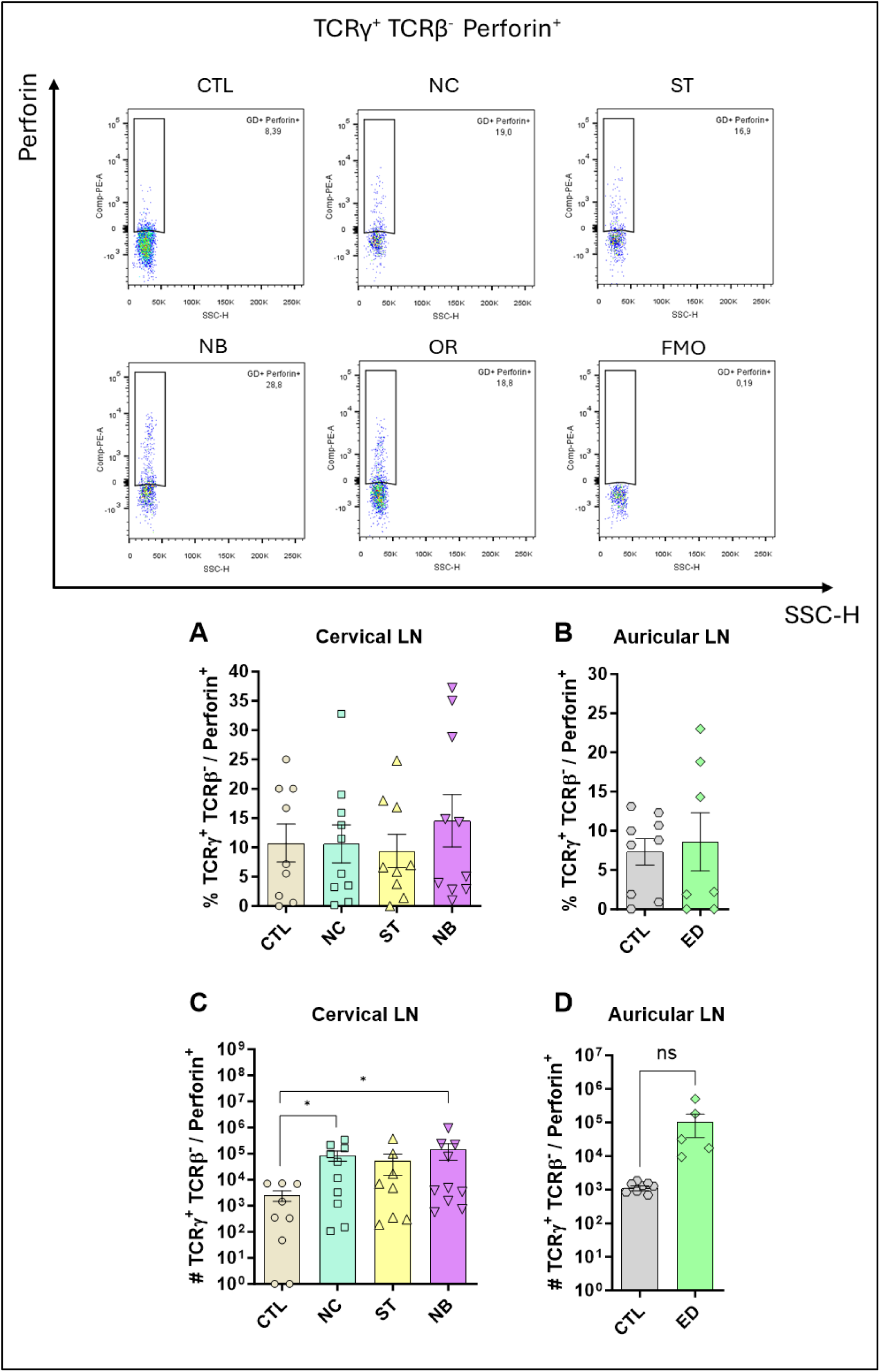
TCRγ^+^ TCRβ^-^ Perforin^+^ profile. The figure shows the representatives and graphs with the percentage of cells in the nasal and ear sites (A) and (B) respectively. In (C) and (D) data on the total number of cells are represented. Data accumulative of two independent experiments. Statistics: plot with Standard Error of The Mean (SEM), t-test was used for all groups and samples p<0,05, **<0,005., 3-6 animals per group in each.

**Figure S26.**
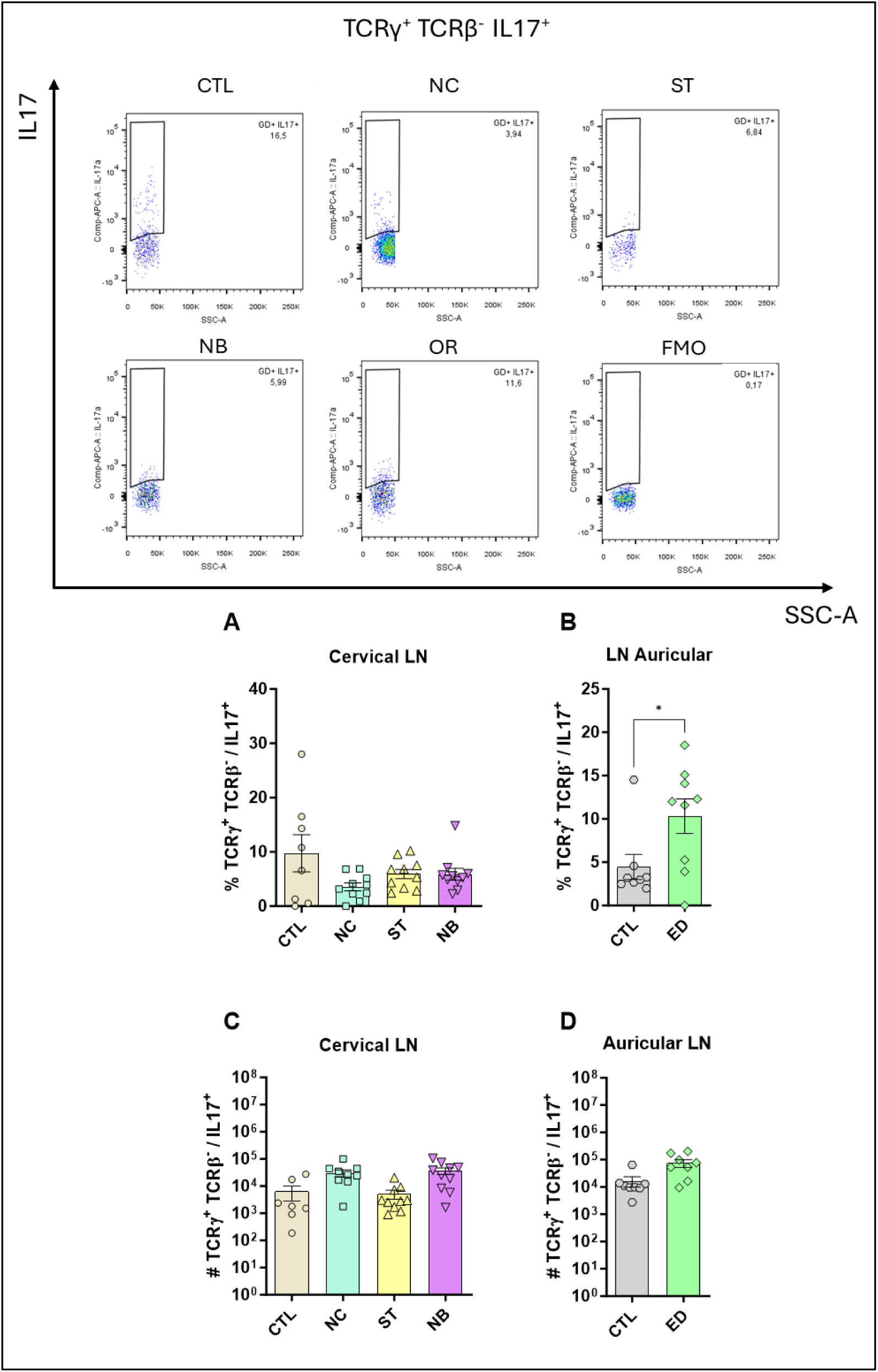
TCRγ^+^ TCRβ^-^ IL-17^+^ profile. The figure shows the representatives and graphs with the percentage of cells in the nasal and ear sites (A) and (B) respectively. In (C) and (D) data on the total number of cells are represented. Data accumulative of two independent experiments. Statistics: plot with Standard Error of The Mean (SEM), t-test was used for all groups and samples *p<0,05, **<0,005., 3-6 animals per group in each.

**Figure S27.**
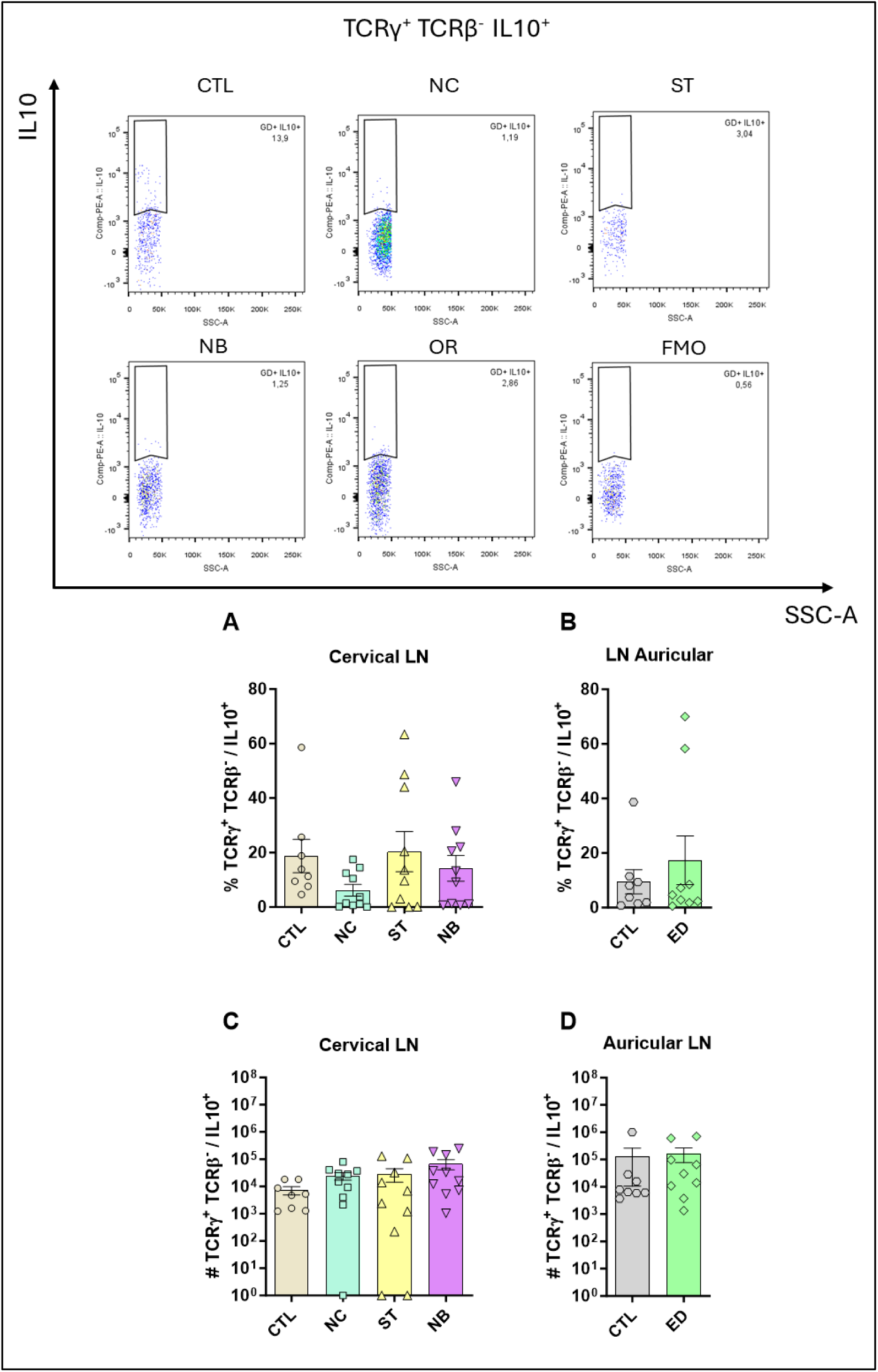
TCRγ^+^ TCRβ^-^ IL-10^+^ profile. The figure shows the representatives and graphs with the percentage of cells in the nasal and ear sites (A) and (B) respectively. In (C) and (D) data on the total number of cells are represented. Data accumulative of two independent experiments. Statistics: plot with Standard Error of The Mean (SEM), t-test was used for all groups and samples *p<0,05, **<0,005., 3-6 animals per group in each.

**Table S1.** BD LSR Fortessa X20 Cytometer Lasers and Filters Configuration.

| **Laser** | **Filter**<br>**BP** | **Filter**<br>**LP** | **Laser** | **Filter**<br>**BP** | **Filter**<br>**LP** |
| --- | --- | --- | --- | --- | --- |
| **Violet** | 780/60 | 750 | **Yellow-**<br>**Green** | 780/60 | 750 |
|  | 710/50 | 690 |  | 710/50 | 685 |
|  | 670/30 | 655 |  | 670/30 | 635 |
|  | 610/20 | 595 |  | 60/20 | 600 |
|  | 525/50 | 505 |  | 586/15 | --- |
|  | 450/50 | --- |  |  |  |
| **Blue** | 710/50 | 690 | **Red** | 780/60 | 750 |
|  | 530/30 | 505 |  | 730/45 | 710 |
|  |  |  |  | 670/30 | --- |

**Table S2.** Main staining antibodies markers, colors and concentrations.

| Extracellular |  |  | Intracellular |  |  |
| --- | --- | --- | --- | --- | --- |
| Antibody | Fluorochrome | Concentration | Antibody | Fluorochrome | Concentration |
| anti-CD3 | BV605 | 1µg/mL | anti-IL-10 | PE | 2µg/mL |
| anti-CD4 | BV785 | 1µg/mL | anti-IL-17 | APC | 2µg/mL |
| anti-CD8 | Percp Cy5.5 | 1µg/mL | anti-IFN-γ | FITC | 2µg/mL |
| anti-TCRβ | BV650 | 1µg/mL | anti-Perforin | PE | 2µg/mL |
| anti-TCRγ | BV510 | 1µg/mL | anti-Granzyme b | APC | 2µg/mL |
| anti-CD25 | PE Cy7 | 1µg/mL | anti-CD107a | APC Cy7 | 2µg/mL |
| anti-PD-1 | PE-Cy7 | 1µg/mL | anti-FoxP3 | AF488 | 2µg/mL |

